# Butyrate as a growth factor of *Clostridium acetobutylicum*

**DOI:** 10.1101/2024.07.15.603595

**Authors:** Hyeongmin Seo, Sofia H. Capece, John D. Hill, Jonathan K. Otten, Eleftherios T. Papoutsakis

## Abstract

The butyrate biosynthetic pathway not only contributes to electron management and energy generation in butyrate forming bacteria, but also confers evolutionary advantages to the host by inhibiting the growth of surrounding butyrate-sensitive microbes. Proteomic data suggest that butyrate may lead to lysine butyrylation, a lesser-known post-translational modification, which might affect enzyme catalysis and thus cellular metabolism. Although high levels of butyrate induce toxic stress responses, it is not known if butyrate at non-toxic levels influences cellular processes such as growth, health, metabolism, and sporulation. Here, we show that butyrate stimulates cellular processes of *Clostridium acetobutylicum*, a model butyrate forming Firmicute. First, we deleted the 3-hydroxybutyryl-CoA dehydrogenase gene (*hbd*) from the *C. acetobutylicum* chromosome in order to eliminate the butyrate synthetic pathway and thus butyrate formation. For rapid genome engineering, a xylose inducible Cas9 cassette was chromosomally integrated and utilized for the one-step markerless gene deletions. The addition of non-toxic levels of butyrate revealed that butyrate has a profound effect on the growth, health, and sporulation of *C. acetobutylicum*. By further deleting *spo0A*, the gene of the master regulator of sporulation, and followed by butyrate addition experiments, we conclude that butyrate affects cellular metabolism through both Spo0A dependent and independent mechanisms. We also deleted the *hbd* gene from the chromosome of the asporogenous *C. acetobutylicum* M5 strain lacking the pSOL1 plasmid to examine the potential involvement of pSOL1 genes on the observed butyrate effects. Addition of the precursor of butyrate biosynthesis crotonate to the *hbd* deficient M5 strain was used to probe the role of butyrate biosynthesis pathway in electron and metabolic fluxes. Finally, we found that butyrate addition can enhance the growth of the non-butyrate forming *Clostridium saccharolyticum*. Our data suggest that butyrate functions as a stimulator of cellular processes, like a growth factor, in *C. acetobutylicum* and other *Clostridium* organisms, and may thus be as a modulator of microbial population dynamics.

**Highlights:**

- Deployed chromosomally integrated spCas9 for markerless one-step *Clostridium acetobutylicum* genome engineering.
- Deleted 3-hydroxybutyryl-CoA dehydrogenase gene (*hbd*) from *Clostridium acetobutylicum* to elucidate the roles of butyrate in cellular processes.
- Demonstrated butyrate as a growth factor stimulating cellular processes in *Clostridium acetobutylicum* and potentially other *Clostridium* species.
- Suggested butyrate as a potential modulator of microbial population based on different responses of microbes against butyrate.

## 1. Introduction

Butyrate (butyric acid) is a four-carbon short-chain fatty acid naturally produced by anaerobic microbes. Butyrate producing bacteria are attracting significant interest across the food, medical, and chemical industries due to the multifaceted benefits of butyrate, including promoting a healthy gut microbiome and colon, as well as its potential applications in diverse chemical and biofuel production processes (Canani et al., 2011; Tracy et al., 2012). Butyrate promotes the growth and/or differentiation of mammalian cells, frequently acting as a growth factor for cultured cells both *in vitro* and *in vivo* (Li and Li, 2006; Prasad and Sinha, 1976), by contributing to histone acetylation that globally regulates gene expression (Candido et al., 1978; Gerhauser, 2018). Furthermore, the importance of butyrate in gastrointestinal health has been extensively examined over the past two decades (Canani et al., 2011; Hodgkinson et al., 2023). In contrast to mammalian cells, mechanisms by which butyrate may influence the physiological processes of microorganisms remain largely unexplored. For biotechnological applications, aside from the stress effects of high butyrate concentrations (Alsaker and Papoutsakis, 2005; Venkataramanan et al., 2013; Venkataramanan et al., 2015), of interest are the effects of butyrate on the cellular behaviors of butyrate forming bacteria, notably on clostridia, either in pure cultures or synthetic consortia (Benomar et al., 2015; Charubin et al., 2018; Charubin et al., 2020a; Charubin and Papoutsakis, 2019).

Belonging to phylum Firmicutes (Bacillota), clostridia are major butyrate producers in microbial communities (Mountzouris et al., 2002). *Clostridium acetobutylicum*, which was historically employed for the industrial acetone-butanol-ethanol (ABE) fermentation, serves as a model strain for understanding clostridial sporulation (Al-Hinai et al., 2015; Jones et al., 2008), transition from acidogenesis to solventogenesis (Amador-Noguez et al., 2011; Durre et al., 1995; Jones et al., 2018), regulatory networks (Paredes et al., 2004; Wang et al., 2013), and more broadly the physiology of solventogenic butyrate forming clostridia (Paredes et al., 2004; Wang et al., 2013).

The butyrate biosynthetic pathway plays an important role in the physiology of butyrate forming clostridia. Especially important is the bifurcating reduction of crotonyl-CoA to butyryl-CoA for electron management and dephosphorylation of butyryl-phosphate to butyrate for ATP generation (Figs. 1A, B). Furthermore, the intermediate metabolite butyryl-phosphate was suggested as a key phosphoryl group donor to phosphorylate and thus activate Spo0A, the highly preserved master regulator of sporulation in Firmicutes (Liu et al., 2003; Paredes et al., 2005; Ravagnani et al., 2000; Steiner et al., 2011; Zhao et al., 2005). As such, butyrate biosynthesis in *C. acetobutylicum* has been hypothesized to be closely related to several regulatory processes including solventogenesis program (Alsaker et al., 2004; Husemann and Papoutsakis, 1988). Of particular interest regarding butyrate’s role in the metabolic regulation in *C. acetobutylicum* are post-translational modifications (PTMs). Identification of global site-specific PTMs in *C. acetobutylicum* suggested that butyrate or a precursor and/or derivative metabolite could play an important role in various protein PTMs (Xu et al., 2018a; Xu et al., 2018b). Given the importance of butyrate forming clostridia in various microbiomes, the role of butyrate can be more complex than what is currently known.

**Figure 1.**
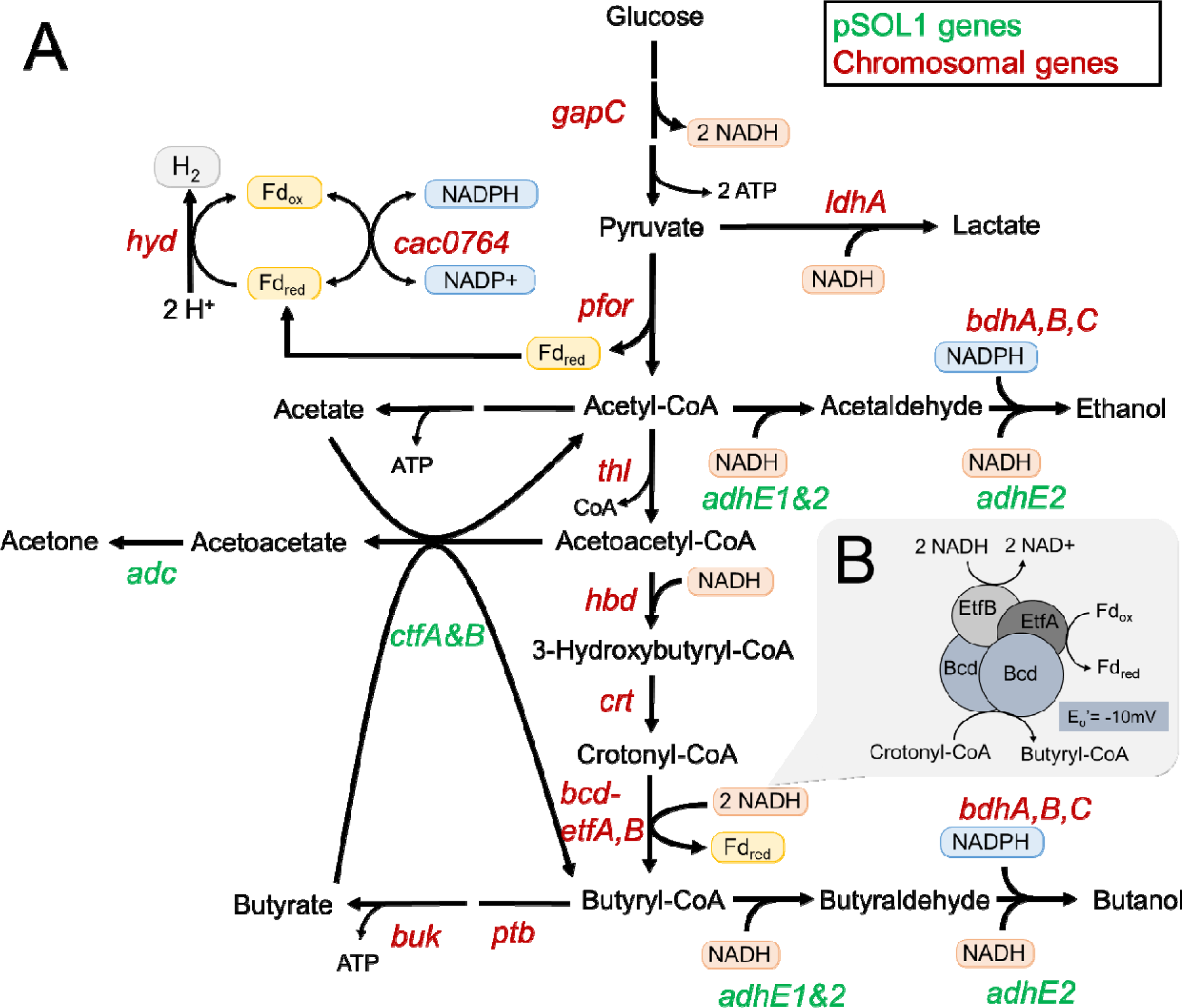
The primary metabolism of *C. acetobutylicum* related to this study. (A) The central metabolic pathway. Green colored genes are encoded in the pSOL1 megaplasmid, and red colored genes are encoded in the chromosome. (B) Details of the bifurcating crotonyl-CoA reduction by the Bcd-EtfAB complex within the butyrate biosynthetic pathway. Gene names and abbreviations: *hyd*, hydrogenase; *gapC*, Glyceraldehyde 3-phosphate dehydrogenase; *ldhA*, lactate dehydrogenase; *pfor*, pyruvate:ferredoxin oxidoreductase; *bdhA, B, C*; butanol dehydrogenases units A,B,C; *thl*, thiolase; *adc*, acetoacetate decarboxylase; *ctfA, B*, acetoacetyl-CoA:acetate/butyrate CoA transferases, units A and B; *hbd*, 3-hydroxybutyryl-CoA dehydrogenase; *crt*, crotonase; *bcd*, butyryl-CoA dehydrogenase; *etfA, B*, electron transferring flavoproteins, units A and B; *ptb*, phosphate butyryltransferase; *buk*, butyrate kinase; *adhE1, 2*, bifunctional aldehyde/alcohol dehydrogenase 1 and 2; Fd_ox_, oxidized ferredoxin; Fd_red_, reduced ferredoxin.

In this study, using genetic modifications and metabolite complementation, we demonstrate the importance of butyrate on *C. acetobutylicum* cell growth and health. Two *C. acetobutylicum* strains were studied: the wild-type (WT) ATCC824 strain containing the pSOL1 plasmid and its M5 derivative lacking the pSOL1 plasmid (Cornillot et al., 1997). The 192 kb size pSOL1 plasmid contains several genes contributing to cellular processes such as solvent production, electron management, and sporulation (Fig. 1A). Through clean deletion of 3-hydroxybutyryl-CoA dehydrogenase (*hbd*) gene from chromosome, we eliminated butyrate formation from the two *C. acetobutylicum* strains. Addition of butyrate and its precursor crotonate to the engineered strains revealed that butyrate is important for robust cell growth as well as sporulation. We further discovered that butyrate enhanced biomass accumulation of a non-butyrate forming *C. saccharolyticum*. While butyrate is not a substrate for cell growth, it promotes cell growth and the developmental program of sporulation. Therefore, we propose that butyrate acts as a growth factor for *C. acetobutylicum* and potentially other clostridia.

## 2. Materials and Methods

### 2.1 Media and cell cultivation

#### 2.1.1 Media composition

The Luria-Bertani (LB) medium was used for propagation of *Escherichia coli.* For the cell growth characterization of *E. coli* in the presence of butyrate, a semi-defined M9 medium consisting of 10 g/L glucose, 6.8 g/L sodium phosphate dibasic, 3.0 g/L potassium phosphate monobasic, 0.5 g/L sodium chloride, 1.0 g/L yeast extract, 1.0 g/L ammonium chloride, 0.12 g/L magnesium sulfate, 0.01 g/L calcium chloride, and 0.4 mg/L ferrous chloride was used. For *C. acetobutylicum* genetic engineering, 2xYTG medium consisting of 16 g/L tryptone, 10 g/L yeast extract, 5 g/L sodium chloride was used. For engineering the *hbd* deficient strains, 30 mM sodium butyrate was added to the 2xYTG medium (2xYTGB). All the *C. acetobutylicum* strains were characterized in clostridial growth medium (CGM) containing 60 g/L glucose, 5 g/L yeast extract, 4 g/L sodium acetate, 2 g/L ammonium sulfate, 2 g/L asparagine, 1 g/L sodium chloride, 0.75 g/L monobasic potassium phosphate, 0.75 g/L dibasic potassium phosphate, 0.7 g/L magnesium sulfate heptahydrate, 17 mg/L manganese sulfate heptahydrate, 10 mg/L iron sulfate heptahydrate, 4 mg/L 4-aminobenzoic acid (Lee et al., 2012). For solid plates, 15 g/L agar were added to the media. For CRISPR/Cas9 genome editing, the CGM agar medium containing 50 g/L xylose instead of glucose (CGMX) was used (Wilding-Steele et al., 2021). Sodium butyrate or crotonic acid was added to the culture media as needed, and pH was adjusted to 6.5 using 5 M sodium hydroxide solution before autoclaving. For some of *C. acetobutylicum* characterization and *C. ljungdahlii* DSM 13528 (ATCC 55383) cell growth, Turbo-CGM (T-CGM) containing 1 g/L monobasic potassium phosphate, 1.25 g/L dibasic potassium phosphate, 1 g/L sodium chloride, 10 mg/L manganese sulfate heptahydrate, 0.348 g/L magnesium sulfate heptahydrate, 10 mg/L iron sulfate heptahydrate, 2 g/L asparagine, 5 g/L yeast extract, 2 g/L ammonium sulfate, 2.46 g/L sodium acetate, 0.2 mg/L sodium tungstate dihydrate, 20 mg/L calcium chloride dihydrate, and 4 mg/L of 4-aminobenzoic acid was used (Charubin and Papoutsakis, 2019). *C. saccharolyticum* WM1 (ATCC 35040) cell growth was characterized in a modified Turbo-CGM medium (T-CGM-CC). All media were deoxygenized by incubating for at least 24 hours inside an anaerobic chamber (Thermo Forma 1025, Thermo Fisher Scientific, MA, USA) after autoclaving. Media were supplemented with the appropriate antibiotics at the following concentrations: 50 μg/mL kanamycin, 30 μg/mL chloramphenicol, 100 μg/mL ampicillin, 100 μg/mL erythromycin, and 10 μg/mL thiamphenicol.

#### 2.1.2 C. acetobutylicum culture for characterization

A single colony on an agar plate was inoculated in 4 mL 2xYTG or 2xYTGB. For sporulating strains, the inoculated cells were heat shocked at 80. The cells were incubated inside an anaerobic chamber at 37. Once grew up, the cells were transferred to 10 mL of a fresh CGM or CGMB, and the optical cell density was measured using a GENESYS 150 UV-Vis spectrophotometer at 600nm wavelength (Thermo Fisher Scientific, MA, USA).

#### 2.1.3 C. saccharolyticum culture for butyrate sensitivity test

For media containing butyrate, 30 mM sodium butyrate was added to the T-CGM-CC before autoclaving. The medium was deoxygenated in an anaerobic chamber under atmospheric blend of 85% N_2_, 10% CO_2_, and 5% H_2_. To initiate the cell culture, a 1.3 mL frozen glycerol stock of *C. saccharolyticum* was inoculated in 80 mL of T-CGM-CC with the cap loose to allow for gas exchange. After being left to grow overnight, the pH of the media was increased to 6.8 to let the cells grow further before bioreactor inoculation. For pH controlled *C. saccharolyticum* culture, small 150 mL scale bioreactors were employed as previously described (Otten et al., 2022). To maintain anaerobic condition, nitrogen gas was continuously fed into the bioreactors at the flow rate of 5 mL/min. The pH was controlled with Bluelab pH Controllers, which dispense boluses of sterile 1.8 sodium hydroxide. *C. saccharolyticum* was added into deoxygenized T-CGM-CC to a target OD_600nm_ of 0.1-0.2.

#### 2.1.4 C. ljungdahlii culture for butyrate sensitivity test

A glycerol frozen stock of *C. ljungdahlii* was inoculated in T-CGM containing 5 g/L fructose but no glucose and incubated at 37 for 2-3 days in an anaerobic chamber. 0.5 mL of the cells were inoculated in a 160 mL serum bottle containing 20 mL of T-CGM with 30 mM butyrate or without butyrate. The serum bottle was capped with a rubber cap and sealed with an aluminum seal inside the chamber and incubated at 37 and 90 rpm in a shaking incubator outside the chamber.

#### 2.1.5 Escherichia coli culture for butyrate sensitivity

A single colony of *E. coli* DH5α was inoculated in LB medium and aerobically incubated at 37 and 200 rpm overnight. 0.5 mL of the cells were inoculated in a 160 mL of serum bottle containing 20 mL of the semi-defined M9 medium with 30 mM butyrate or without butyrate, inside an anaerobic chamber. After capping and sealing the bottle with a rubber cap and an aluminum seal inside the chamber, the cells were anaerobically incubated at 37 and 200 rpm in a shaking incubator.

### 2.2 Strain and plasmid construction

#### 2.2.1 Plasmid construction

The bacterial strains and plasmids used in this study are listed in Table 1. The list of primers used in this study is presented in Table S1. All plasmids were constructed using NEBuilder HiFi DNA assembly (New England Biolabs, MA, USA). The PCR fragments were amplified using Phusion DNA polymerase or AccuPrime DNA polymerase (Thermo Fisher Scientific, MA, USA). Successful construction of the plasmids was confirmed by colony PCR using Phire DNA polymerase (Thermo Fisher Scientific, MA, USA), and Sanger sequencing.

**Table 1.**
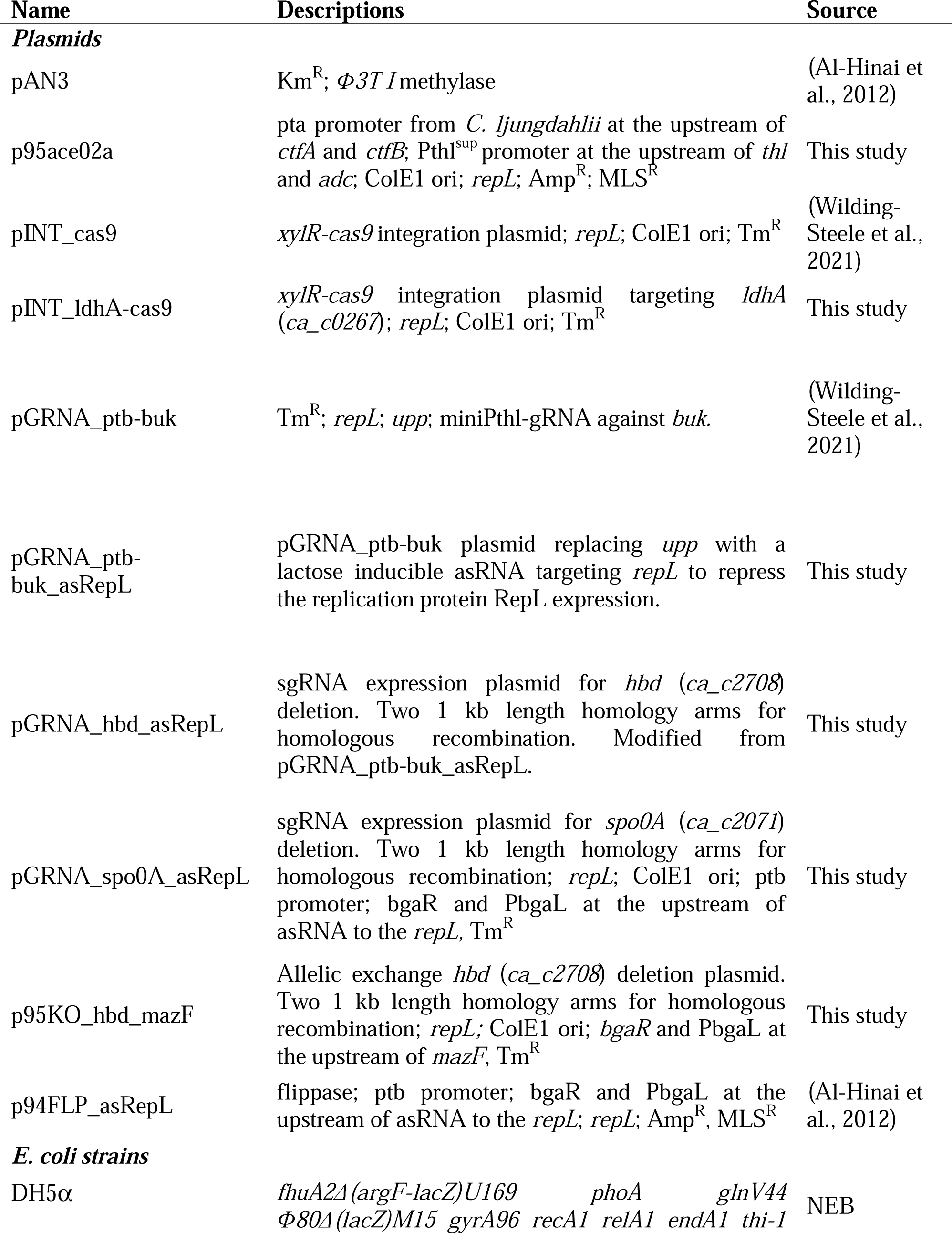
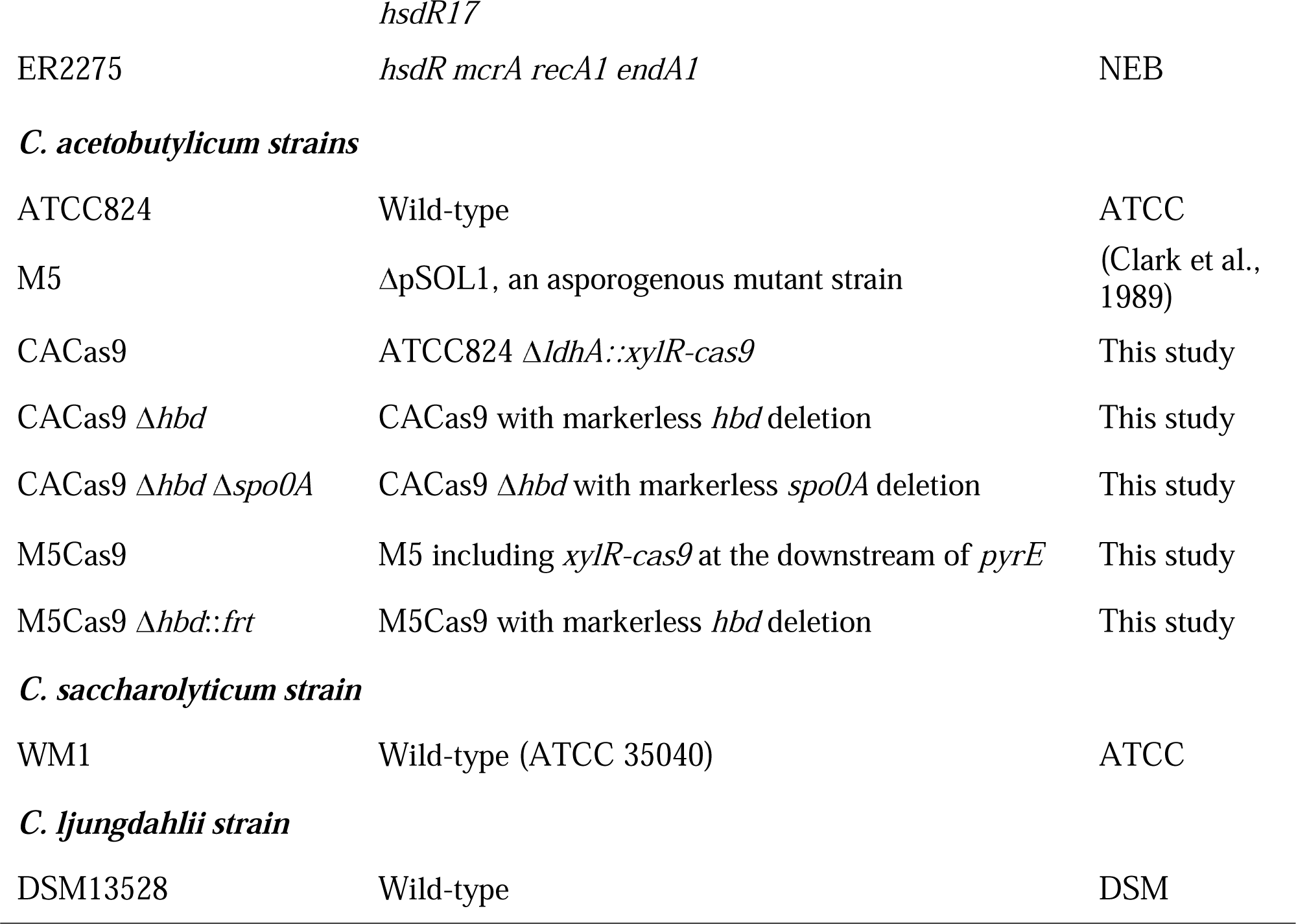
A list of bacterial strains and plasmids used in this study.

#### 2.2.2 Transformation of C. acetobutylicum

Electroporation of *C. acetobutylicum* was performed as previously described (Mermelstein and Papoutsakis, 1993). All plasmids were methylated in *E. coli* ER2275 carrying the pAN3 plasmid and electroporated into *C. acetobutylicum*. All procedures were performed inside an anaerobic chamber. The 60 mL culture in 2xYTG at early exponential phase (OD_600nm_ of 0.4-0.8) was harvested by centrifugation at 6,500 × g for 5 minutes. The pellets were washed in 10 mL of ice-cold electroporation buffer (EPB) comprising of 270 mM sucrose in 5 mM sodium phosphate buffer at pH 7.4. After discarding the supernatant, the pellets were concentrated in 2 mL ice-cold EPB. 0.5 mL of the resuspended cells were transferred to a 4 mm electroporation cuvette, followed by addition of 1-5 μg of the methylated DNA plasmids to the cells. After 3 minutes of incubation in ice, a 2.0 kV exponential pulse at 25 µF capacitance and infinite resistance using the Gene Pulser (Bio-Rad, Hercules, CA, USA). Then, the cells were immediately transferred to 4 mL of 2xYTG medium without antibiotics and incubated for 4-6 hours at 37 for recovery. For the *hbd* deficient strains, 2xYTGB was used for the recovery. Then, the cells were harvested by centrifugation 6,500 × g for 5 minutes and plated on a 2xYTG or 2xYTGB agar plate containing erythromycin or thiamphenicol. The cells were incubated at 37 for 3-5 days until transformant colonies appeared.

#### 2.2.3 Construction of CACas9 and M5Cas9

The xylose inducible Cas9 cassette was chromosomally integrated into *C. acetobutylicum* ATCC824 and M5 using pINT_ldhA-cas9 and pINT_cas9 (Wilding-Steele et al., 2021), respectively. For the chromosomal integration, a single transformant colony was inoculated in 2xYTG containing thiamphenicol and incubated until the cell growth reached an early exponential phase (OD_600nm_ of 0.4-0.6). 100 μL of the cell culture were plated on CGMX agar medium containing thiamphenicol and incubated at 37 for 2-4 days until colonies appeared. Individual colonies were diagnosed by PCR for successful integration of the Cas9 cassette at the intended chromosomal location. The constructs were finally confirmed by Sanger sequencing of the PCR product. Confirmed colonies were passaged 4-5 times on 2xYTG agar medium without thiamphenicol through streaking for plasmid curing. Finally, the colonies without growth in the presence of thiamphenicol were tested by PCR again for confirmation of the integration of the Cas9 cassette.

#### 2.2.4 Genome editing of CACas9

Plasmid pGRNA_asRepL was constructed by inserting a lactose-inducible antisense RNA against RepL (asRepL) to pGRNA_bukptb plasmid (Table 1) and used as the backbone plasmid for sgRNA and homology arm cloning. asRepL accelerates plasmid curing by repressing the replication protein expression in the presence of lactose (Al-Hinai et al., 2012). Guide RNA sequences were designed using CHOPCHOP (https://chopchop.cbu.uib.no/) (Shen et al., 2018), and a gene fragment including promoter sequences, the 20 bp length guide RNA, scaffold, and terminator were synthesized (Integrated DNA Technologies, IA, USA). The gene fragment was assembled with PCR amplified homology arm sequences to construct a sgRNA plasmid. Gene deletion was performed by inducing the chromosomal Cas9 expression by plating on CGMX (Wilding-Steele et al., 2021). After the successful deletion, plasmid curing was performed on 2xYTG or 2xYTGB agar plates containing 50 mM lactose.

### 2.3 Analytical methods

#### 2.3.1 Quantitative reverse transcription polymerase chain reaction (RT-qPCR)

RT-qPCR was performed to quantify the relative expression levels of two representative solventogenic genes in *C. acetobutylicum* (i.e., *ctfA*, *adhE1*). The *fabZ* was used as a housekeeping control. CACas9 Δ*hbd* cells cultured in CGM or CGMB were harvested at the early stationary phase, followed by cell disruption and RNA extraction using TRIzol (Thermo Fisher Scientific, MA, USA) and RNeasy Kit (Qiagen, Germany). The RNA samples were quantified and used as templates for one-step RT-qPCR using Luna Universal One-Step RT-qPCR kit (NEB, MA, USA). Real time PCR amplification was performed using CFX96 touch real-time PCR detection system (Biorad, CA, USA). After determining amplification efficiencies of the primers, *fabZ* relative fold expression was quantified using Pfaffl method (Pfaffl, 2001).

#### 2.3.2 Sporulation assay

Individual colonies of CACas9 Δ*hbd* were randomly selected and mixed via vortexing in 1 mL of T-CGM inside an anaerobic chamber. 500 µL of the cell suspension was inoculated into 4.5 mL of T-CGM with or without 50 mM butyrate and anaerobically incubated at 37 for 120 hours and transferred into a refrigerator (4) outside the anaerobic chamber for 8 days to promote sporulation. Then, 1 mL of the culture was pelleted and resuspended in 150 µL of sterile phosphate buffered saline (1xPBS, pH 7.4), followed by serial 20x dilutions in 1xPBS. The serially diluted cultures were incubated in a heat block at 80 for 10 minutes. 100 µL of each dilution was spread on deoxygenated 2xYTG plates containing 37 mM butyrate butyrate and incubated anaerobically for 2 days and colonies were then counted. For the butyrate containing condition, plates with between 50-200 colonies were used to estimate frequency. For the non-butyrate condition, only one or no colony appeared even on the least diluted plates. Therefore, we assumed a spore frequency of 1.5 spores per mL, which represents the detection limit for this assay.

#### 2.3.3 Microscopic examination of endospore formation

A single colony of CACas9 Δ*hbd* strain on 2xYTG without butyrate were heat-shocked at 80 for 10 minutes and incubated at 37 for 3-5 days until early exponential phase (OD_600nm_ of 0.4-0.8). Then, 15-20 µL of the liquid culture was streaked onto 2xYTG (pH 5.8) plates with or without butyrate. The strains were allowed to grow for 6 days. Then, a smear from each strain was made on a glass microscope slide and stained with malachite green and safranin (Schaeffer-Fulton spore stain kit, Sigma-Aldrich, MO, USA) to visualize endospores. Stained cells were examined using a Zeiss Axioplan 2 Upright Light Microscope under the bright field setting with a 63x objective with oil immersion. Images captured with the Axiocam 512 Color Camera were used to count the endospores through ImageJ cell counter plug-in.

#### 2.3.4 High-performance liquid chromatography (HPLC) analysis

An HPLC system (Agilent, CA, USA) was used to measure metabolite and sugar concentrations. 700 μL of cell culture supernatant was filtered through a 0.2-micron filter and run with into 5 mM H_2_SO_4_ at 0.5 mL/min on an Aminex HPX-87H column (Biorad Inc., CA, USA) at 15. The metabolites were detected and quantified using a refractive index detector.

## 3. Results

### 3.1 Inactivation of the butyrate-formation pathway in *C. acetobutylicum* necessitates the use of exogenous butyrate for robust cell growth

Deletion of *hbd* can eliminate butyrate formation by disrupting the butyrate biosynthetic pathway at the first pathway step below the thiolase reaction (Fig. 1A). For rapid and clean gene deletion, we employed a tightly controlled xylose inducible Cas9 genome editing tool (Wilding-Steele et al., 2021). By integrating the large size of Cas9 cassette into the chromosome, this system enables sequential one-step gene deletion using a relatively small plasmid that expresses a designed sgRNA (Wilding-Steele et al., 2021). The lactate dehydrogenase gene (*ldhA, ca_c0267*) was chosen as the integration location for the Cas9 cassette, aiming to attenuate the undesirable lactate and 2-hydroxyvalerate formation (Yoo et al., 2017). The WT *C. acetobutylicum* strain harboring the chromosomally integrated Cas9 cassette was successfully constructed and designated as CACas9 (Fig. 2A). Since only miniscule amounts of lactate are produced under typical *C. acetobutylicum* culture conditions, the chromosomal integration of Cas9 cassette at the *ldhA* location did not change the growth and solvent formation characteristics of the engineered strain when compared to the parent strain (Supplementary Figure 1). By introducing a plasmid harboring an sgRNA targeting *hbd* with 1 kb homology arms (pGRNA_hbd_asRepL; see Methods) and inducing expression of the chromosomal Cas9, the *hbd* gene was successfully deleted (Fig. 2B). The resulting strain, CACas9 Δ*hbd*, exhibited relatively smaller colony size compared to the parental CACas9 on the 2xYTG plate during the *hbd* deletion process. Compared to CACas9, CACas9 Δ*hbd* exhibited a longer lag phase (up to 5 days vs. 1 day) following the standard heat-shock spore germination protocol. Furthermore, CACas9 Δ*hbd* exhibited poor cell growth, reaching a maximum cell density (OD_600nm_) of 2.9 ± 1.9 after 94 hours of culture in CGM liquid medium (Fig. 2C). No detectable butyrate or butanol was produced, indicating that the deletion of *hbd* disrupted the butyrate biosynthetic pathway. Ethanol and acetate were produced as the two major products, reaching concentrations of 60 ± 41 mM and 38 ± 23 mM, respectively. Only 0.16 ± 0.14 mM of acetone were formed, indicating weak solventogenic activities. CACas9 Δ*hbd* exhibited relatively high variations in cell growth and metabolite formation across the biological replicates, suggesting that the deletion of *hbd* significantly compromised cell robustness.

**Figure 2.**
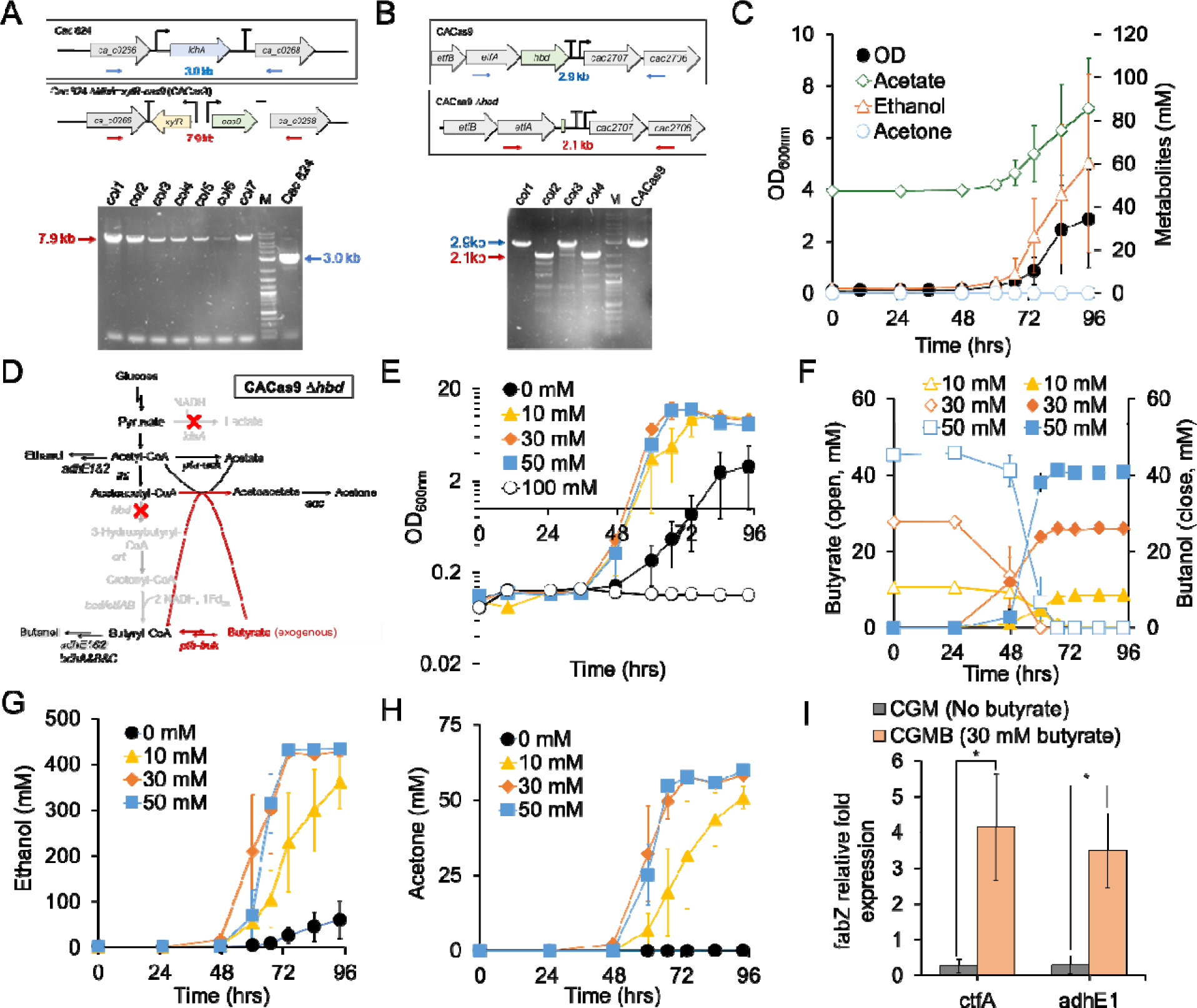
Deletion of the butyrate biosynthetic pathway in *C. acetobutylicum* ATCC824. (A) Integration of the xylose inducible Cas9 cassette into the chromosome of *C. acetobutylicum* ATCC824. A scheme and DNA gel picture of markerless deletion of *hbd* from the chromosome of CACas9. Red and blue arrows indicate the expected band size for successful gene deletion and parental genotype, respectively. col1-7 represent individual colonies screened by colony PCR. M, DNA marker. (B) Deletion of 3-hydroxybutyryl-CoA dehydrogenase (*hbd*) gene from the chromosome of CACas9. The red and blue arrows indicate the expected band size for successful gene deletion and parental genotype, respectively. col1-4 represent individual colonies screened by PCR. M, DNA marker. (C) Cell growth and metabolite profiles of CACas9 Δ*hbd* in CGM medium. (D) A scheme of central metabolism of CACas9 Δ*hbd* upon butyrate addition. The red ‘X’ marks indicate the deleted metabolic reactions through corresponding gene deletions. (E-I) Time dependent profiles of CACas9 Δ*hbd* in CGM with different butyrate concentrations. (E) Profiles of cell growth. (F) Butyrate and butanol. (G) Ethanol, (H) Acetone, (I) Relative fold-change in solventogenic gene expressions in CACas9 Δ*hbd*. *, p-value <0.05 from a two-tailed t-test. All the data represent means ± standard deviation from three biological replicates.

Butyryl-phosphate, a butyrate derivative metabolite, has been suggested to induce acetone and butanol production in *C. acetobutylicum* (Harris et al., 2000; Husemann and Papoutsakis, 1988; Zhao et al., 2005). Deletion of *hbd* eliminated butyrate formation from glucose that would prevent the formation of intracellular butyryl-phosphate for the induction of solventogenesis. We thus hypothesized that adding butyrate to CACas9 Δ*hbd* would restore vigorous cell growth and acetone formation. Because CACas9 Δ*hbd* retains the two potential pathways for butyrate uptake, exogenous butyrate could be assimilated by the cells (Fig. 2D). To test this hypothesis, we compared growth and metabolite profiles of CACas9 Δ*hbd* in CGM containing butyrate at different concentrations (Fig. 2E-H). Addition of butyrate as low as 10 mM significantly improved CACas9 Δ*hbd* growth with 12 hours shorter lag phase compared to no butyrate addition (Fig. 2E). With 30 mM butyrate, the maximum cell density (OD_600nm_) was 12.0 ± 2.4, 4.1-fold higher than the cell density without butyrate (2.9 ± 1.9). These results clearly indicate that addition of butyrate restored robust cell growth. No cell growth was observed at 100 mM of butyrate, likely due to butyrate toxicity. The added butyrate was assimilated and converted to butanol at the conversion rates of 90-94% (mol/mol) before the cell growth reached stationary phase (Fig. 2F). When more than 10 mM butyrate were added, CACas9 Δ*hbd* produced ethanol up to 431 mM, corresponding to the yield of 1.3 (mol ethanol per mol glucose) (Fig. 2G). Acetone was produced up to 60 mM (Fig. 2H), suggesting that solventogenesis was significantly restored by addition of butyrate. Most of the acetone was produced after 60 h, when butyrate had already been converted to butanol. Therefore, acetone formation was accompanied by acetate assimilation rather than butyrate assimilation (Figs. 2F, H, and Supplementary Figure 2A). Interestingly, cell growth, glucose consumption, and metabolite formation in the presence of 10 mM butyrate were relatively slower compared to 30 mM and 50 mM butyrate (Figs. 2E, G, H, and Supplementary Figure 2B), indicating that 10 mM butyrate were not sufficient to fully restore cell growth and metabolism. CACas9 Δ*hbd* growth rates in a modified CGM containing 30 mM sodium butyrate (CGMB) reached 0.36 ± 0.03 h^-1^, 2.5-fold higher than the growth rates in CGM (0.14 ± 0.01 h^-1^) (Supplementary Figure 2C). We further hypothesized that, in CACas9 Δ*hbd*, expression of core genes needed for solvent production (*sol* operon) would not be induced without butyrate. The *sol* operon consists of *adhE1* (*aad*), *ctfA*, and *ctfB* genes (Fischer et al., 1993; Nair et al., 1994). AdhE1 is a bifunctional aldehyde-alcohol dehydrogenase, and CtfA&B are the two units of the acetoacetyl-CoA:acetate/butyrate:CoA transferase which is responsible for the first step in the acetone formation pathway that generates acetoacetyl-CoA (Fig. 1). Expression of the *sol* operon is activated by phosphorylated Spo0A (Spo0A-P) (Thormann et al., 2002). As a proxy to the presence of activated Spo0A-P (Jones et al., 2008), expression levels of two representative *sol* operon genes, *ctfA* and *adhE1*, with and without butyrate were examined via RT-qPCR (Fig. 2I). The addition of 30 mM butyrate significantly induced expression of *ctfA* and *adhE1* leading to 28- and 22-fold higher expression levels, respectively, compared to without butyrate. These results suggested that defective Spo0A activation in the absence of butyrate attenuated solventogenic gene expression and acetone formation (Harris et al., 2002).

### 3.2 Exogenous butyrate is critical for sporulation of the *hbd* deficient *C. acetobutylicum*, further supporting its role in activating Spo0A

Binding of phosphorylated Spo0A to its DNA binding motif directly regulates expression of more than 200 genes in *C. acetobutylicum*. Additionally, it indirectly regulates expression of many more genes through the Spo0A regulated sigma factors, notably SigF, SigG, and SigK (Al-Hinai et al., 2015; Jones et al., 2008; Paredes et al., 2004). This regulation controls various cellular programs including carbohydrate metabolism, electron transport, motility, chemotaxis, energy production, and signaling proteins (Alsaker et al., 2004; Tomas et al., 2003). Spo0A-P controls many other metabolic processes and gene expression as well as sporulation in *C. acetobutylicum* (Fig. 3A). To further examine the hypothesis that exogenous butyrate significantly affects Spo0A phosphorylation, which is necessary for robust sporulation, we measured the sporulation frequency as a proxy for Spo0A activation. We compared the sporulation frequency of CACas9 Δ*hbd* under the culture conditions without butyrate and with butyrate (Fig. 3B). The sporulation frequency of CACas9 Δ*hbd* increased by the three orders of magnitude (2,650-fold) when butyrate was added to the medium (Fig. 3C). The sporulation frequency of CACas9 Δ*hbd* was 1.9 ± 0.8 CFU/mL, suggesting that the sporulation program of CACas9 Δ*hbd* was highly attenuated by the disruption of the butyrate biosynthesis pathway. The impact of the exogenous butyrate on the sporulation program was also confirmed by microscopy (Fig. 3D, E, and Supplementary Figure 3). In the absence of butyrate, only one cell out of 1,133 cells displayed sporulation characteristics detected by endospore staining. In the presence of butyrate, a significantly greater number of cells exhibited endospore formation. Based on the exceptionally high conservation of the Spo0A protein sequence in *Bacillus* and *Clostridium* organisms, and its canonical DNA motif, the 0A box (‘-TGNCGAA-3’) (Ravagnani et al., 2000), unphosphorylated Spo0A still binds to the 0A boxes with different affinities that leads to different developmental consequences (Baldus et al., 1994; Fujita et al., 2005). Thus, the marginal sporulation frequency was to be expected even if Spo0A was not fully phosphorylated.

**Figure 3.**
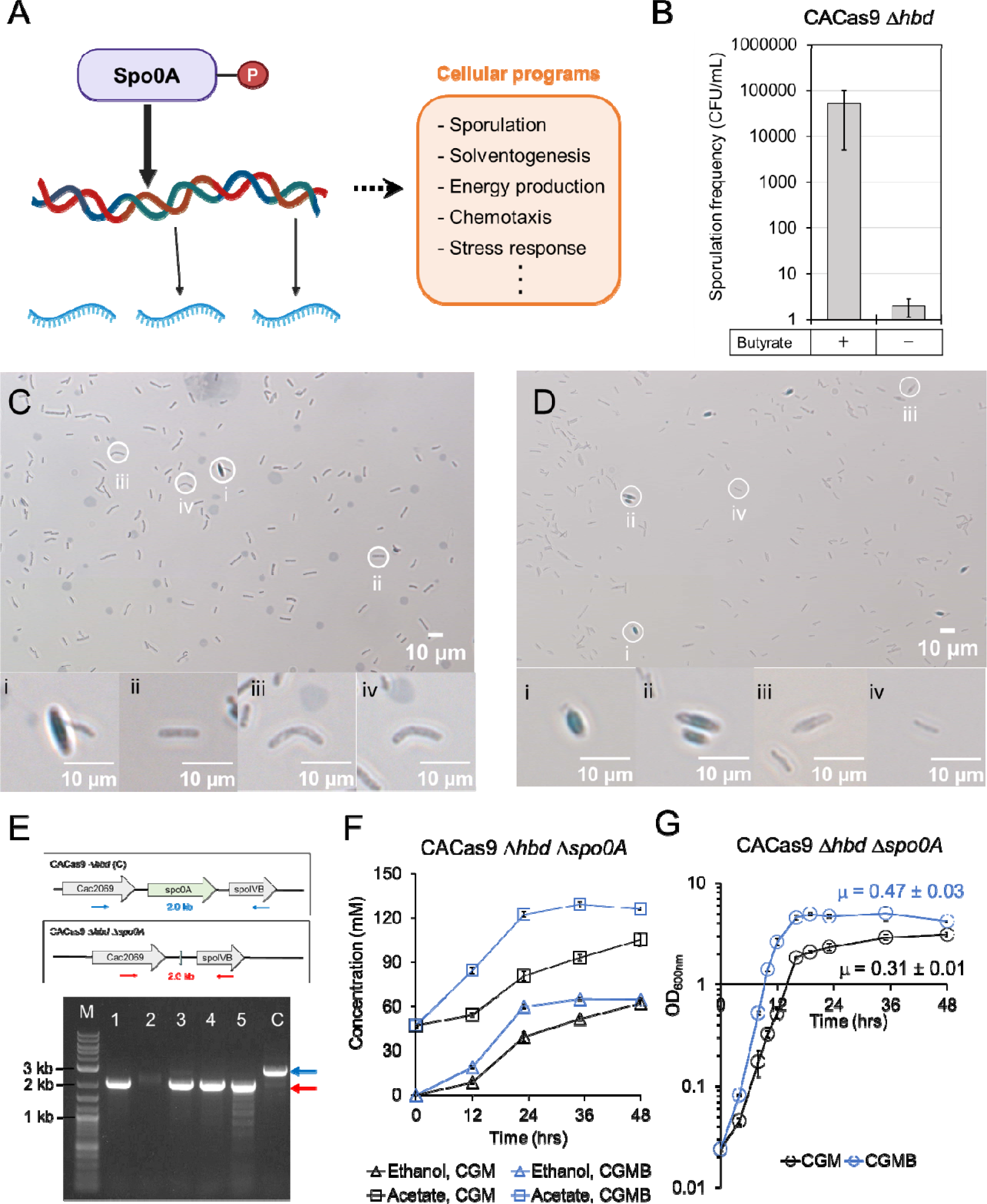
Effects of butyrate addition on Spo0A controlled phenotypes. (A) A scheme of Spo0A mediated regulation of cellular processes. (B) Sporulation assay of CACas9 Δ*hbd* with and without 50 mM butyrate addition. (C) Light microscopic examination of CACas9 Δ*hbd* cells without butyrate addition. i-iv, individual cells presented in the enlarged picture. (D) Light microscopic examination of CACas9 Δ*hbd* cells with 30 mM butyrate addition. i-iv, individual cells presented in the enlarged picture. (E) Deletion of *spo0A* from the chromosome of CACas9 Δ*hbd*. 1-5 represent individual colonies diagnosed by colony PCR. M, DNA marker; C, CACas9 Δ*hbd*. The red and blue arrows indicate the expected band size for successful gene deletion and parental genotype, respectively. (F) Profiles of acetate and ethanol production by CACas9 Δ*hbd* Δ*spo0A* in CGM and CGMB media. (G) Cell growth of CACas9 Δ*hbd* Δ*spo0A* in CGM and CGMB media. All the data represent mean ± standard deviation from three biological replicates.

### 3.3 Activation of Spo0A is only partially responsible for the butyrate effects

Due to the global regulator role of Spo0A, defective Spo0A phosphorylation could impact the growth and metabolism of CACas9 Δ*hbd* in various ways. To further probe the relationship between butyrate and Spo0A, we deleted *spo0A* from the chromosome of CACas9 Δ*hbd* (Fig. 3E) and examined the impact of butyrate compared to that on CACas9 Δ*hbd*. As expected, CACas9 Δ*hbd* Δ*spo0A* did not sporulate and as such could not survive the standard heat shock process. Without butyrate, the specific growth rate of CACas9 Δ*hbd* Δ*spo0A* (no *spo0A* expression) (0.31 ± 0.01 h^-1^; Fig. 3F) was 2.2-fold higher than that of CACas9 Δ*hbd* (Spo0A activation is deficient) (0.14 ± 0.01 h^-1^; Fig. 2D). The somewhat unexpected results suggest that even without activation, Spo0A affected cell growth and metabolism in complex ways, notably negatively affecting the apparent growth rates. This was not observed in the *C. acetobutylicum* WT strain upon spo0A inactivation (Harris et al., 2002), suggesting that impact of spo0A deletion is context dependent. In *Bacillus subtilis*, Spo0A exerts a negative impact on DNA replication by binding on the origin of replication (Castilla-Llorente et al., 2006). CACas9 Δ*hbd* Δ*spo0A* showed acetate and ethanol as the major products (Fig. 3F), with a small amount of acetone formation (Supplementary Figure 4). The specific growth rate (0.47 ± 0.03 h^-1^) and maximum cell density (OD_600nm_ = 5.03 ± 0.78) of CACas9 Δ*hbd* Δ*spo0A* with butyrate addition were 53% and 61% higher, respectively, than those without butyrate addition (0.31 ± 0.01 h^-1^ and OD_600nm_ = 3.11 ± 0.14) (Fig. 3G). While the cell growth enhancement by butyrate was substantial, it was not as pronounced as the case of CACas9Δ*hbd* (i.e., 2.5- and 7.2-fold enhancement of specific growth rate and maximum OD_600nm_, respectively) (Fig. 2D). This suggests that while butyrate likely affected cell growth and metabolism through Spo0A phosphorylation (strain CACas9 Δ*hbd*), its impact on cell growth, health, and metabolism was also significant in the absence of Spo0A expression (strain CACas9 Δ*hbd* Δ*spo0A*). One possible explanation of this observation is that butyryl-phosphate is a phosphoryl group donor for phosphorylating not only Spo0A, but also other regulatory proteins (such as AbrB protein) and enzymes through the large number of two component kinase systems as well as orphan kinases (Paredes et al., 2005). The impact of such phosphorylation events in controlling growth and metabolism is well established (Garcia-Garcia et al., 2016).

### 3.4 Exogenous butyrate enhances cell growth of asporogenous *C. acetobutylicum* M5 strain lacking the butyrate biosynthetic pathway

We next questioned if the butyrate effects are related to the role of pSOL1 megaplasmid, which encodes 178 genes, including all essential genes for acetone and butanol formation (the *sol* operon), as well as genes related to the hydrogenase assembly, the sporulation program, electron management machinery, and alternative carbon-substrate utilization, among others (Nolling et al., 2001). Thus, we tested the non-solventogenic, asporogenous *C. acetobutylicum* M5 strain, which lacks the pSOL1 megaplasmid and produces butyrate as the major product due to the loss of key pSOL1 genes responsible for acetone and butanol production (Fig. 1A). *C. acetobutylicum* M5 produces a small amount of ethanol as at least two alcohol dehydrogenases are encoded on the chromosome (Nair and Papoutsakis, 1994; Petersen et al., 1991; Welch et al., 1989). To test the impact of butyrate on *C. acetobutylicum* M5, we deleted the *hbd* gene from its chromosome.

First, we tried to use the same Cas9 approach as for the WT strain. We chromosomally integrated a xylose inducible Cas9 cassette at the location between hydrogenase (*hydA*) and orotate phosphoribosyltransferase (*pyrE*) genes (Supplementary Figure 5), a site previously used for chromosomal gene integrations (Ehsaan et al., 2016; Wilding-Steele et al., 2021). Unfortunately, deleting the *hbd* gene from the M5Cas9 strain using the Cas9 was unsuccessful. Then, we tried a different approach using the two-step allelic exchange method using a thiamphenicol resistance gene (*cat*) and the *mazF* toxin gene as the selection and counterselection markers, respectively (Al-Hinai et al., 2012). Using the two-step method, an *hbd* deficient *C. acetobutylicum* M5Cas9 strain, namely M5Cas9 Δ*hbd*::*cat*, was successfully constructed and isolated (Fig. 4A). Compared to the parental M5Cas9 strain, M5Cas9 Δ*hbd*::*cat* exhibited a longer lag phase (up to 4 days vs. 1 day) and significantly reduced maximum OD_600nm_ (less than 0.5 vs. 7.2) that inhibited effective subsequent transformation with p94FLP_asRepL plasmid required for the excision of *cat* selectable marker through FRT recombination (Al-Hinai et al., 2012). Adaptation of the M5Cas9 Δ*hbd*::*cat* strain to the liquid culture medium significantly improved cell growth achieving maximum OD_600nm_ up to 1.5 (Supplementary Figure 6). Following that, *cat* was successfully excised via FRT-FLP recombination (Fig. 4B). A strain with the markerless *hbd* deletion was isolated and named as M5Cas9 Δ*hbd* (Fig. 4C). This strain produced lactate as the major product, and no butyrate was formed as expected (Fig. 4D).

**Figure 4.**
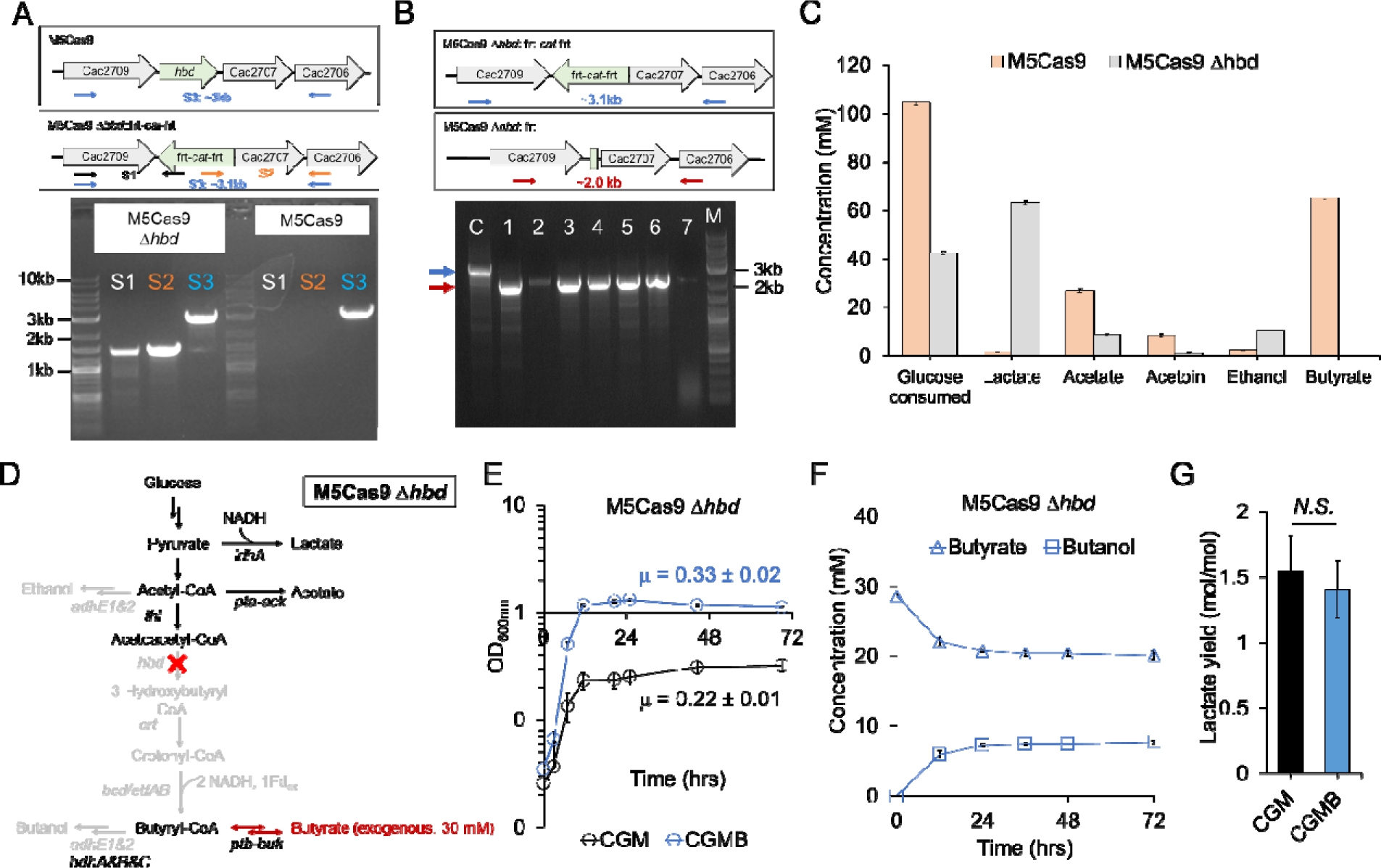
Deletion of butyrate biosynthetic pathway in *C. acetobutylicum* M5 strain. (A) Deletion of the 3-hydroxybutyryl-CoA dehydrogenase (*hbd*) gene from the chromosome of the asporogenous *C. acetobutylicum* M5 strain (M5Cas9). The S1, S2, and S3 indicate the combination of primers used to amplify the different sites to confirm the deletion of *hbd*. (B) Excision of the *cat* selectable marker in M5Cas9 Δ*hbd::frt-cat-frt* via FRT-FLP recombination. 1-7 represent individual colony screened by colony PCR. M, DNA marker; C, M5Cas9 Δ*hbd::frt-cat-frt*. Red and blue arrows indicate the expected band size for successful FRT-FLP recombination and parental genotype, respectively. (C) The metabolite profile of M5Cas9 and M5Cas9 Δ*hbd* after 48 hours of culture in CGM medium. (D) A scheme of the central metabolism of M5Cas9 Δ*hbd* strain upon butyrate addition. The red ‘X’ mark indicates the deleted metabolic reaction through the *hbd* deletion. (E) Cell growth of M5Cas9 Δ*hbd* in CGM and CGMB media. (F) Profiles of butyrate and butanol of M5Cas9 Δ*hbd* culture in CGMB medium. (G) Lactate yields in CGM and CGMB media. All data represent mean ± standard deviation from three biological replicates.

To test the effects of exogenous butyrate on the cell growth of M5Cas9 Δ*hbd*, growth rates of M5Cas9 Δ*hbd* in CGM and CGMB media were measured (Fig. 4E). Butyrate addition significantly enhanced cell growth rates of M5Cas9 Δ*hbd* by 33%. The maximum OD_600nm_ in the presence of 30 mM butyrate was 4-fold higher (1.32 ± 0.04) than the maximum OD_600nm_ in the absence of butyrate (0.32 ± 0.01). This substantial cell growth enhancement was consistent with what was observed with the two CACas9 Δ*hbd* strains (Figs. 2D, 3F), further supporting the beneficial effects of butyrate on the *C. acetobutylicum* cell growth. Because M5Cas9 lacks the acetoacetyl-CoA:acetate/butyrate transferase genes (*ctfA/B*), the only possible pathway for butyrate uptake is the reversible phosphotransbutyrylase and butyrate kinase (Ptb-Buk) reaction that consume 1 mol of ATP per 1 mol of butyrate uptaken (Fig. 1A). Surprisingly, 7 mM out of the 30 mM added butyrate were converted into butanol by M5Cas9 Δ*hbd* (Fig. 4F), which lacks the pSOL1 plasmid that encodes two major aldehyde/alcohol dehydrogenases *adhE1* and *adhE2* responsible for the conversion of butyryl-CoA to butyraldehyde and butanol (Fontaine et al., 2002; Nair et al., 1994). The partial conversion of butyrate to butanol suggests that the *C. acetobutylicum* chromosome codes for a weak butyraldehyde dehydrogenase activity, potentially by the chromosomal alcohol dehydrogenases BdhA, B, and C (Walter et al., 1992; Yoo et al., 2015). The yields of lactate were not statistically different in the presence of butyrate (Fig. 4G), indicating that the beneficial effects of butyrate were not significantly related to metabolism in M5Cas9 Δ*hbd*. Phosphorylation of Spo0A and other cellular proteins remain the logical explanation as for the CACas9 Δ*hbd* strains above.

### 3.5 Assimilation of crotonate improves cell growth and alters electron-dependent metabolic flux of the *hbd* deficient M5 strain

Butyrate producing microbes synthesize crotonyl-CoA as a conserved intermediate in all butyrate biosynthetic pathways (Vital et al., 2014). Reduction of crotonyl-CoA to butyryl-CoA involves electron bifurcating reaction catalyzed by the complex of the butyryl-CoA dehydrogenase (coded by the *bcd* gene) with the electron-transferring flavoprotein (coded by the *etfA/B* genes) (Herrmann et al., 2008) (Fig. 1B), an important energy conservation route in butyrate producing anaerobic bacteria (Peters et al., 2016). Motivated by the strong metabolic flux and cell growth changes caused by *hbd* gene deletion from the chromosome of M5Cas9 Δ*hbd*, we examined if crotonate addition to generate crotonyl-CoA would activate the bifurcating crotonyl-CoA reduction reaction. The *hbd* gene is located at the end of a 5 kb length polycistronic butyryl-CoA synthesis (BCS) operon consisting of five genes, *crt-bcd-etfB-etfA-hbd* (Fig. 1). It was unclear whether deleting the *hbd* gene would destabilize the polycistronic mRNA, potentially suppressing the expression of the remaining BCS genes. Testing the impact of crotonate addition would address the uncertainty as well. Then, the challenge was identifying a mechanism for crotonate uptake.

Although crotonate could be potentially assimilated by promiscuous Ptb-Buk activity (Fischer et al., 2010), addition of 30 mM crotonate did not enhance the growth of M5Cas9 Δ*hbd*, and there was no apparent crotonate uptake and butyrate/butanol formation (Supplementary Figure 7). This finding suggested that the chromosomal Ptb-Buk activities were not sufficient for converting crotonate to crotonyl-CoA. Another potential mechanism was using the acetoacetyl-CoA:acetate/butyrate CoA transferases (CoAT), coded by *ctfA/B* in pSOL1 plasmid (Fig. 1). The native CtfAB has been shown to exhibit a small activity against crotonate (Wiesenborn et al., 1989). To enable a continuous operation of the CoAT activity given that it generates acetoacetate (Fig. 1), the full acetone pathway was required to be expressed. Thus, we transformed M5Cas9 Δ*hbd* with p95ace02a that harbors four acetone biosynthesis genes (*thl*, thiolase; *ctfA/B*, CoAT; *adc*, acetoacetate decarboxylase) under the control of strong constitutive promoters (Ppta_clj_, and Pthl_sup_) (Fig. 5A) (Charubin et al., 2020b; Streett et al., 2019). Then, 20, 50, and 100 mM of croconate were added to cultures, and the growth of M5Cas9 Δ*hbd*/p95ace02a was monitored and compared to control (no crotonate addition). There was significantly improved cell growth of M5Cas9 Δ*hbd*/p95ace02a by the addition of crotonate (Fig. 5B). The maximum OD_600nm_ from the culture with 50-100 mM crotonate reached up to 2.87 ± 0.21, which was 7-fold higher compared to the culture without crotonate (OD_600nm_ = 0.44 ± 0.14). The specific cell growth rates in the presence of 20, 50, 100 mM crotonate were 0.28 ± 0.02 h^-1^, 0.34 ± 0.01 h^-1^, and 0.30 ± 0.01 h^-1^, respectively. In contrast, the specific cell growth rates without crotonate were only 0.10 ± 0.02 (h^-1^). As hypothesized, crotonate was assimilated into the cell through the CoAT reaction, producing acetone (Figs. 5C, D). Crotonate uptake resulted in significant formation of equimolar butyrate and butanol (Figs. 5E, F). Furthermore, the electron carrying metabolic fluxes of lactate and H_2_ formation were altered by the crotonate addition, indicating that the crotonyl-CoA reduction changed the electron flux from lactate formation to H_2_ formation (Figs. 5G, H). The strong correlation between crotonate uptake and H_2_ formation suggest that the bifurcating ferredoxin reduction from the butyrate biosynthetic pathway is a key contributor to H_2_ production in *C. acetobutylicum* (Figs. 5C, H).

**Figure 5.**
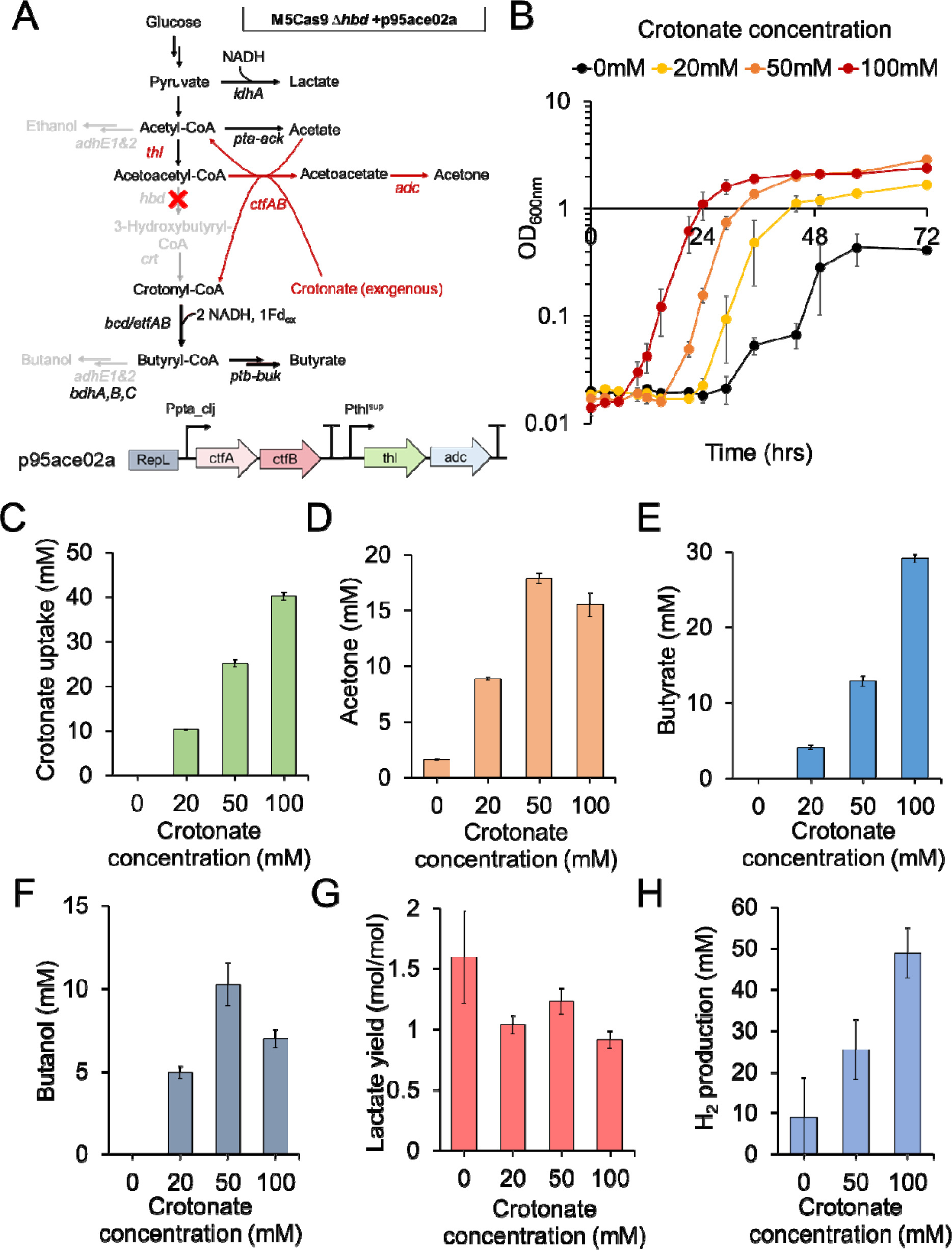
Effects of crotonate addition to cultures of M5Cas9 Δ*hbd* transformed with the p95ace02a plasmid. (A) A map of central metabolism of M5Cas9 Δ*hbd* with p95ace02a (M5Cas9 Δ*hbd*/p95ace02a), and the operon architecture of p95ace02a. The red arrows indicate the overexpressed reactions by p95ace02a. (B) Cell growth profiles of M5Cas9 Δ*hbd*/p95ace02a at the different concentrations of added crotonate. (C) The amount of consumed crotonate after 72 hours at the different concentrations of added crotonate. (D-F) Concentrations of produced metabolites after 72 hours at the different concentrations of added crotonate. (D) Acetone. (E) Butyrate. (F) Butanol. (G) Lactate yields after 72 hours at the different concentrations of added crotonate. (H) Concentration of produced H_2_ (calculated based on liquid culture volume) at the different concentrations of added crotonate. All data represent mean ± standard deviation from three biological replicates.

### 3.6 Exploring butyrate effects on non-butyrate forming microbes

Given the beneficial effects of butyrate on *C. acetobutylicum*, we further questioned whether butyrate also modulates the growth of non-butyrate forming microbes. We investigated the effects of butyrate addition on the growth of *C. saccharolyticum*, *C. ljungdahlii*, and *Escherichia coli*, which do not naturally synthesize butyrate. These microbes coexist with butyrate forming bacteria in natural microbiome and synthetic co-cultures (Benomar et al., 2015; Otten et al., 2022).

Addition of 30 mM butyrate enhanced the growth of *C. saccharolyticum*, which produces acetate and ethanol as the main metabolites (Fig. 6A). The maximum OD_600nm_ was increased by 40% from 1.85 ± 0.04 to 2.63 ± 0.08. In addition, acetate titers were improved by 60% in the presence of 30 mM butyrate (Fig. 6B). The titers of the other major metabolite, ethanol, were not significantly altered. The results suggest that the exogenous butyrate changed the metabolic flux of *C. saccharolyticum*, particularly at the acetyl-CoA node. There was no apparent butyrate consumption or butanol formation by *C. saccharolyticum*. The improved cell growth of *C. saccharolyticum* was further confirmed under various culture conditions (Figs. 6C, D), suggesting that butyrate played a key role in improved biomass accumulation. In contrast, the addition of 30 mM butyrate did not significantly affect the growth of *C. ljungdahlii,* and rather inhibited the growth of *E. coli* (Supplementary Figure 8). The results indicated that the non-butyrate forming microbes exhibit different and variable responses to butyrate, with some being inhibited while others exhibiting stimulated growth.

**Figure 6.**
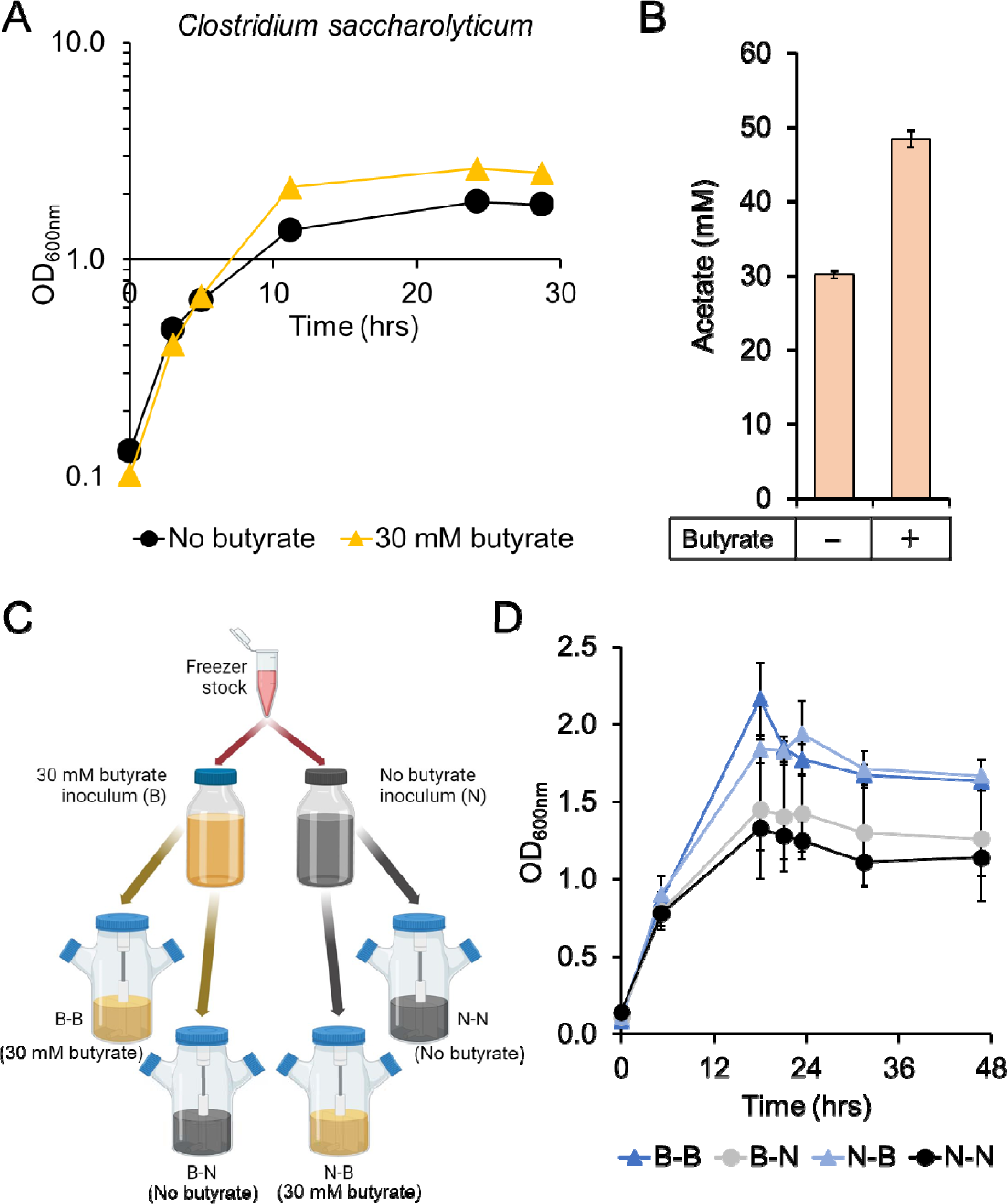
Effects of 30 mM added butyrate in *C. saccharolyticum* culture. (A) Cell growth in T-CGM-CC medium with or without butyrate. The data represent mean ± standard deviation from four biological replicates. (B) Acetate titers after 30 hours of the culture. The data represent mean ± standard deviation from four biological replicates. (C) A scheme of inoculum preparation and small bioreactor operation. (D) Cell growth under different preculture and main culture conditions per panel C. The data represent mean ± standard deviation from two biological replicates.

## 4. Discussion

*C. acetobutylicum* has been employed as a powerful microbial host for metabolic engineering and synthetic biology applications over the past decades. However, the biological importance of butyrate biosynthesis in *C. acetobutylicum* has not been fully explored. In this study, we demonstrated that butyrate stimulates the growth and health of *C. acetobutylicum* through butyrate addition in three genetically engineered strains, CACas9 Δ*hbd* (Fig. 2D), CACas9 Δ*hbd* Δ*spo0A* (Fig. 3F), and M5Cas9 Δ*hbd* (Fig. 4E). The activating impact of butyrate addition supports the idea that butyryl-phosphate may act as a phosphoryl group donor for Spo0A activation (Harris et al., 2000; Zhao et al., 2005).

Several studies have reported different phenotypes of *C. acetobutylicum* strains with engineered butyrate metabolism using different genetic engineering tools (Desai et al., 1999; Green et al., 1996; Lehmann and Lutke-Eversloh, 2011; Nguyen et al., 2018; Zhao et al., 2005). For example, inactivation of the *ptb* gene using an intron-based gene disruption tool ClosTron showed high levels of ethanol and acetone production (Lehmann et al., 2012b), different from other studies which inactivated *ptb* and/or *buk* using allelic exchange (Green et al., 1996; Yoo et al., 2017) or Cas9 (Wilding-Steele et al., 2021). In contrast to the findings reported here, ClosTron-based *hbd* gene inactivation showed insignificant effects on the cell growth of *C. acetobutylicum* despite of undetectable butyrate formation (Lehmann and Lutke-Eversloh, 2011). One possible explanation is that the insertion-based mutagenesis using ClosTron resulted in unknown polar effects, secondary mutations (Lutke-Eversloh, 2014), or a leaky phenotype (Lehmann et al., 2012a). For instance, deletion of *adhE2* from pSOL1 using ClosTron caused a polar effect disrupting the nearby solventogenic operon (Cooksley et al., 2012; Yoo et al., 2016). Clean gene deletion using allelic exchange or Cas9 genome editing employed in this study is assumed to be free from such concerns.

Recent proteomic analyses suggested that the PTMs of lysine (e.g., acetylation and butyrylation) and phosphorylation of proteins takes place at high frequency in *C. acetobutylicum* (Xu et al., 2018a; Xu et al., 2018b). There are 1,078 butyrylated lysine sites among 373 proteins including ribosomal proteins, and metabolic enzymes of glycolysis, butyrate and fatty acid metabolism (Xu et al., 2018b). Butyryl-CoA possibly serves as a butyryl group donor for protein butyrylation including lysine butyrylation of Spo0A, potentially catalyzed by general control non-depressible 5 (Gcn5)-related N-Acetyltransferases (GNATs) (Xu et al., 2018b). The wide distribution of butyrylated proteins suggests that butyrate biosynthesis might contribute to various cellular processes via PTMs. Our results experimentally support the importance of butyrate biosynthesis for facilitating various cellular processes via protein phosphorylation and, possibly, via protein butyrylation in *C. acetobutylicum* (Fig. 7). Phosphorylation of Spo0A is catalyzed by orphan kinases, but it remains uncertain if butyryl-phosphate participates in this phosphorylation process (Harris et al., 2000; Steiner et al., 2011; Zhao et al., 2005).

**Figure 7.**
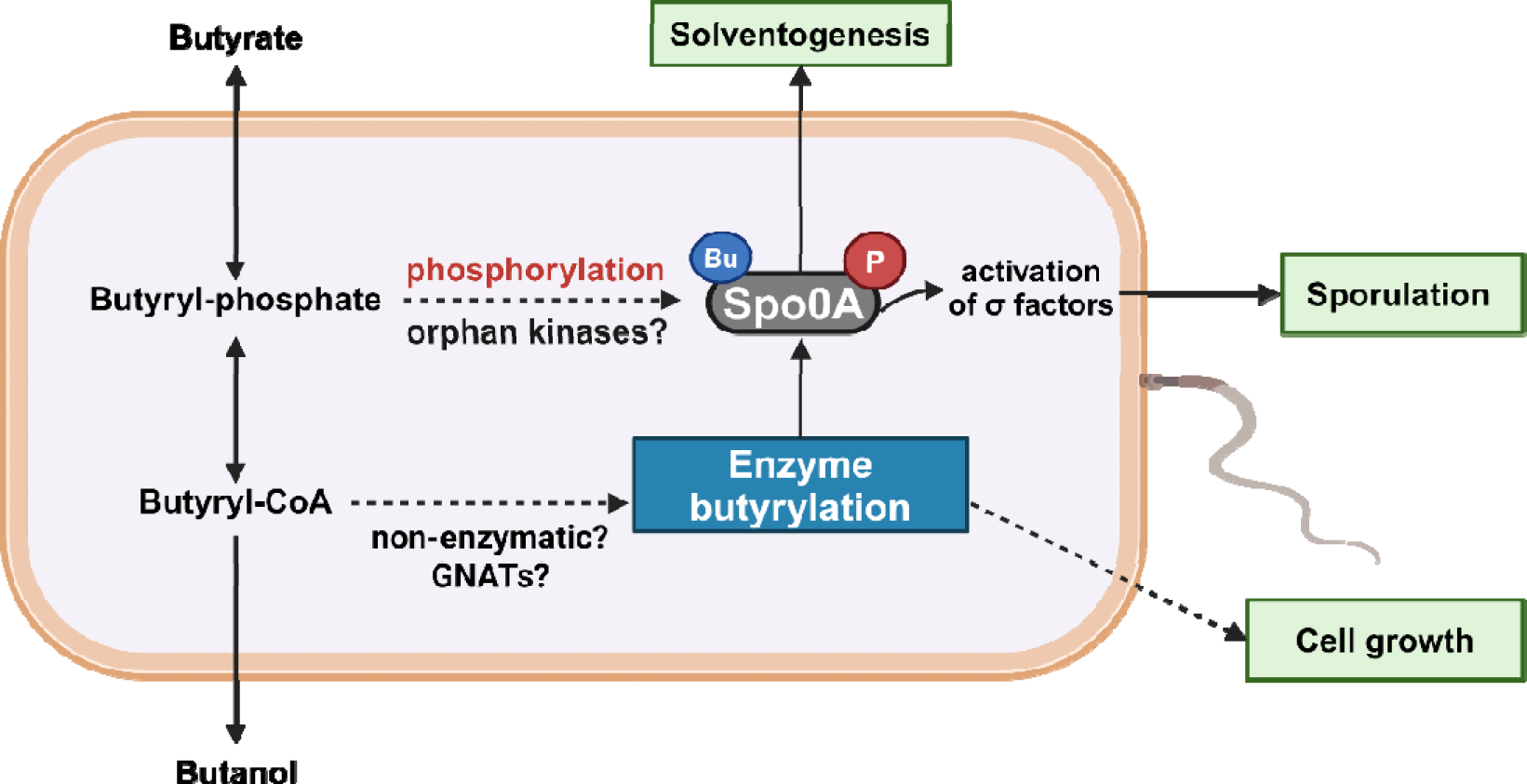
A proposed mechanism of action of butyrate as a growth factor for *C. acetobutylicum*. Abbreviation: GNATs, General control non-depressible 5 (Gcn5)-related N-Acetyltransferases.

Because microbes exhibit significantly different responses to butyrate (Fig. 6, Supplementary Figure 8), butyrate could play a key role in modulating microbial population within microbiota. One mechanism of the responses includes the butyrate associated PTMs as proposed in this study. Future studies with a focus on proteomic and transcriptomic responses of microbiota will help understand the mechanism of the different microbial responses to butyrate. Given the fact that many microbes reside in butyrate containing environment, identifying the impact of butyrate on their protein PTMs and cellular programs will help understand microbial population dynamics. Recent advances in genetic engineering techniques for manipulating microbial communities hold promise for elucidating the biological significance of butyrate.

## 5. Conclusions

Butyrate acts as a growth factor for *C. acetobutylicum* by enabling its robust growth and metabolism but without being necessarily metabolized. The mechanism of action is likely related to complicated regulatory machinery including Spo0A mediated gene expression. We propose butyrate as a potential microbial population modulator because cell growth and health of a wide range of microbes within microbiota could be differently influenced by butyrate.

## CRediT Author Contributions

**Hyeongmin Seo**: Conceptualization; Data curation; Formal analysis; Investigation; Methodology; Validation; Visualization; Writing-original draft; Writing-review & editing, **Sofia H. Capece**: Investigation; Methodology; Validation; Visualization; Writing-original draft, **John D. Hill**: Investigation; Methodology; Validation; Roles/Writing-original draft, **Jonathan K. Otten**: Investigation; Methodology; Visualization; Writing-original draft, **Eleftherios T. Papoutsakis**: Conceptualization; Funding acquisition; Project administration; Resources; Writing-review & editing.

## Acknowledgements

This work was financially supported by the U.S. Department of Energy, Advanced Research Projects Agency-Energy (ARPA-E) ECOSynBio under award number # DE-AR0001505. JDH was supported in part by the U.S. Department of Education GAANN Fellowship under grant P200A210065. Illustrations were created with <u>BioRender.com</u>.

## Notes

### Competing Interest Statement

The authors have declared no competing interest.

