## Supplemental file for "Butyrate as a growth factor of *Clostridium acetobutylicum*"

**Supplementary File S1**

**Butyrate as a growth factor of *Clostridium acetobutylicum***

Hyeongmin Seo, Sofia H. Capece, John D. Hill, Jonathan K. Otten, and Eleftherios T. Papoutsakis^§^

Department of Chemical and Biomolecular Engineering, University of Delaware, Newark, DE, USA

**Table S1.** A list of primers and synthesized DNA fragments used in this study. The underlined sequences are the 20 bp length target specific sequences in the sgRNA.

| Primer name | Primer sequence (5’ to 3’) | Description |
| --- | --- | --- |
| ***p95ace02a construction*** | | |
| Ppta-p95 F | AAACTATTGGTTGGAATGGCGTGAAATGCCTAAGTGAAATATATACATATTATAAC | Ppta_clj promoter assembly with plasmid backbone, forward |
| Ppta R | GACTAATCCTCCTCTCTTAAATTTAACACA | Ppta_clj promoter, reverse |
| ctfA-Ppta F | AATTTAAGAGAGGAGGATTAGTCATGAACTCTAAAATAATTAGATTTGAAAATTTAAG | *ctfAB* assembly with Ppta_clj promoter, forward |
| ctfB-Pthl R | AAATTTCAATACTTTTTGTTAAAAATAAAAATAAGAGTTACCTTAAATGGTAACTC | *ctfAB* assembly with Pthl^sup^ promoter at the downstream, reverse |
| Pthl F | TTTTTAACAAAAAGTATTGAAATTTGGTC | Pthl^sup^ promoter, forward |
| Pthl R | GGATCCTAACCTCCTAAATTTTGATAC | Pthl^sup^ promoter, reverse |
| Thl-pthl F | ATCAAAATTTAGGAGGTTAGGATCCATGAAAGAAGTTGTAATAGCTAGTG | *thl* assembly with Pthl^sup^ promoter, forward |
| Thl-p95 R | CTAGCACTTTTCTAGCAATATTGCT | *thl*, reverse |
| adc-thl F | GCAATATTGCTAGAAAAGTGCTAGAGGAAGGTGACTTTTATGTTAAAGGATGAAG | *adc* assembly with *thl*, forward |
| adc-thl R | TAAGAGTTACCATTTAAGGTAACTCTTACTTAAGATAATCATATATAACTTCAGCTC | *adc* assembly with plasmid backbone, reverse |
| p95 BB F | GAGTTACCTTAAATGGTAACTCTTATTTTTTTAATGTCGACTCATA | Plasmid backbone, forward |
| p95 BB R | CACGCCATTCCAACCAATAG | Plasmid backbone, reverse |
| ***pGRNA_asRepL construction*** | | |
| asRepL-pGRNA F | TAGTGAGTCGTATTACGCGCGCTCCGAGATGAAAAGTATTAGGGCTA | bgaR-PbgaL::asRepL assembly with plasmid backbone, forward |
| asRepL-pGRNA R | CGAAGTAAATAAGTCTAGTGTGTTAGACCACAGTCGACCGACCATG | bgaR-PbgaL::asRepL assembly with plasmid backbone, reverse |
| pGRNA-asRepL F | GAGCGCGCGTAATACG | pGRNA plasmid backbone for asRepL cassette cloning, forward |
| pGRNA-asRepL R | GTCTAACACACTAGACTTATTTACTTCG | pGRNA plasmid backbone for asRepL cassette cloning, reverse |
| ***pINT_ldhA-cas9 construction*** | | |
| ldhA LHA F | GGCCGCCGTCTAGAACTAGTGGATCGATATAATAAACGGAACCTTAAAACCAGG | ldhA left homology arm, forward |
| ldhA LHA R | ATGCAACATTGTATTTAAAATATAAAACTTTGT | ldhA left homology arm, reverse |
| ldhA RHA F | GGTAAAAGACCTAAACTCCAAGG | ldhA right homology arm, forward |
| ldhA RHA R | CACGGATCTGGATCATTACGAATT AGATCAATTGAACCTGCAATCGC | ldhA right homology arm, reverse |
| INT-ldhA F | TATATTTTAAATACAATGTTGCATGGTACCAACTCGAGAATCTAGAAAGAGCTC | xylR-cas9 assembly with ldhA homology arms, forward |
| INT-ldhA R | ACCCTTGGAGTTTAGGTCTTTTACCCGGGTCGACGTGTAAGCTCCTGCAG | xylR-cas9 assembly with ldhA homology arms, reverse |
| pGRNA BB F | AATTCGTAATGATCCAGATCCGTG | pGRNA plasmid backbone for assembly with an sgRNA fragment and homology arms, forward |
| pGRNA BB R | TTTTGTTAAAAAGTAGGGCCCGCGGCC | pGRNA plasmid backbone for assembly with an sgRNA fragment and homology arms, reverse |
| ***pGRNA_hbd construction*** | | |
| hbd LHA F | GGCCGCCGTCTAGAACTAGTGGATCATAGATTTCTTATCAAGTCAAAAACCTCC | hbd left homology arm, forward |
| hbd LHA R | CTGTTTAATGTAAGTTTACAAGAATCCCCATTATCAAATG | hbd left homology arm, reverse |
| hbd RHA F | ATTCTTGTAAACTTACATTAAACAGACCTCCCTAAATTTAAAG | hbd right homology arm, forward |
| hbd RHA R | CACGGATCTGGATCATTACGAATTTTGAATTAACAGCTGTTTTACTTGGAC | hbd right homology arm, reverse |
| ***pGRNA_spo0A construction*** | | |
| spo0A LHA F | AATGTATTAATACATTTAACTTTTAACTCCCCTTTATAAAATATAGAAAT | spo0A left homology arm, forward |
| spo0A LHA R | GATCTGGATCATTACGAATTGGACTTTGGGTAAGAGATTCTA | spo0A left homology arm, reverse |
| spo0A RHA F | CCGTCTAGAACTAGTGGATCGGTATTCCTATCATTGTAAGTCCT | spo0A right homology arm, forward |
| spo0A RHA R | TTTATAAAGGGGAGTTAAAAGTTAAATGTATTAATACATTGGTAAAAGTT | spo0A right homology arm, reverse |
| ***p95KO_hbd_mazF construction*** | | |
| p95-bgaR_mazF F | TTTAGAAAAATAAACAAATAGGGGTTCC | Plasmid backbone for cloning bgaR-mazF cassette, forward |
| p95-bgaR_mazF R | ATCTAGGTGAAGATCCTTTTTGATAATC | Plasmid backbone for cloning bgaR-mazF cassette, reverse |
| bgaR-mazF F | TATCAAAAAGGATCTTCACCTAGATCTCGAGATGAAAAGTATTAGGGCTA | bgaR-mazF cassette, forward |
| bgaR-mazF R | ACCCCTATTTGTTTATTTTTCTAAACCCGATTGACGAATACTATGAAAC | bgaR-mazF cassette, reverse |
| p95KO hbd RHA F | GAATTCGCGGCCGCTAAATTCAATTCATTAAACAGACCTCCCTAAATT | hbd right homology arm assembly with frt-cat-frt fragment, forward |
| p95KO hbd RHA R | GATAATGGAGCAGTAAGAGCTATGGTATTGAATTAACAGCTGTTTTACTTG | hbd right homology arm assembly with frt-cat-frt fragment, reverse |
| p95KO hbd LHA F | GTCACTATGCCGAATTCGCCCTTATAGATTTCTTATCAAGTCAAAAACCTCC | hbd left homology arm assembly with frt-cat-frt fragment, forward |
| p95KO hbd LHA R | AAGGGCGAATTCGTTTAAACCTGCTAAGTTTACAAGAATCCCCATTATC | hbd left homology arm assembly with frt-cat-frt fragment, reverse |
| frt-cat-frt F | GCAGGTTTAAACGAATTCGC | frt-cat-frt fragment, forward |
| frt-cat-frt R | AATTGAATTTAGCGGCCGC | frt-cat-frt fragment, reverse |
| ***Genome modification confirmation*** | | |
| ldhA check F | GGTACAGGAAGTTTTGTTAGAGA | PCR diagnosis for xylR-Cas9 insertion at ldhA location, forward |
| ldhA check R | GTAATTCATTTGAAGCAGCACATA | PCR diagnosis for xylR-Cas9 insertion at ldhA location, reverse |
| Cas9-pyrE check F | GGGCTAAAAATGGAACAATACAAAC | PCR diagnosis for xylR-Cas9 insertion at the downstream of pyrE location, forward |
| Cas9-hydA check R | CTCAACATGCTTATTATGCGG | PCR diagnosis for xylR-Cas9 insertion at the downstream of pyrE location, reverse |
| hbd check F | AGCAGCTATTTTAAGTTTACAATTCTT | PCR diagnosis for hbd deletion, forward |
| hbd check R | CATTGGAATTATTAGGTAAAGGTAAGG | PCR diagnosis for hbd deletion, reverse |
| Spo0A check F | GGGAAAAACCTTTCAAGAGATC | PCR diagnosis for spo0A deletion, forward |
| Spo0A check R | AGAAAAGGTAATAAGTAAAGTAAAATGC | PCR diagnosis for spo0A deletion, reverse |
| CAT IF | ATTTGTGTGATATCCACTTTAACGG | PCR diagnosis for hbd deletion through allelic exchange with frt-cat-frt, forward |
| CAT IR | GATTCCTATTTTTACTATGGGGAAA | PCR diagnosis for hbd deletion through allelic exchange with frt-cat-frt, reverse |
| ***RT-qPCR*** | | |
| adhE1 F | GGACTAGCACTAGAGGCAATA | Amplification efficiency (E): 95.5% |
| adhE1 R | GTTGAAGCGTGAGCCATTT |  |
| ctfA F | ATATAGGCAGCAACCCAGATAC | Amplification efficiency (E): 102.9% |
| ctfA R | CCTGCACGTATTCTTTCCACTA |  |
| fabZ F | TGGTAGACAGAGTTGAAGAAATAGA | Amplification efficiency (E): 94.3% |
| fabZ R | ACCTGGGTAATGACCTTGAAATA |  |
| DNA fragments | Sequence (5’ to 3’) | Description |
| *ldhA* sgRNA | GGCCGCGGGCCCTACTTTTTAACAAAATATATTGATAAAAATAATAATAGTGGGTATAATTAAGTTGTTAGTTGGAATTAACGGAGTGAGTTTTAGAGCTAGAAATAGCAAGTTAAAATAAGGCTAGTCCGTTATCAACTTGAAAAAGTGGCACCGAGTCGGTGCTTTTTTGTCGACGCGGCCGCCGTCTAGAACTAGTGGATC | sgRNA targeting *ldhA*, assembly with pGRNA backbone |
| *hbd* sgRNA | GGCCGCGGGCCCTACTTTTTAACAAAATATATTGATAAAAATAATAATAGTGGGTATAATTAAGTTGTTTATGGGTTCAGGAATTGCTCGTTTTAGAGCTAGAAATAGCAAGTTAAAATAAGGCTAGTCCGTTATCAACTTGAAAAAGTGGCACCGAGTCGGTGCTTTTTTGTCGACGCGGCCGCCGTCTAGAACTAGTGGATC | sgRNA targeting *hbd*, assembly with pGRNA backbone |
| *spo0A* sgRNA | GGCCGCGGGCCCTACTTTTTAACAAAATATATTGATAAAAATAATAATAGTGGGTATAATTAAGTTGTTTTATTATATCGAGAACAACAGTTTTAGAGCTAGAAATAGCAAGTTAAAATAAGGCTAGTCCGTTATCAACTTGAAAAAGTGGCACCGAGTCGGTGCTTTTTTGTCGACGCGGCCGCCGTCTAGAACTAGTGGATC | sgRNA targeting *spo0A*, assembly with pGRNA backbone |

**Supplementary Figure 1.** Comparison of growth and the pH profile of *C. acetobutylicum* ATCC824 (Cac 824) and the CACas9 strain. Rep1, Rep2, Rep3 indicate individual biological replicates. (A) Cell growth of Cac 824. (B) Cell growth of CACas9. (C) pH profile of Cac 824 culture. (D) pH profile of CACas9 culture.

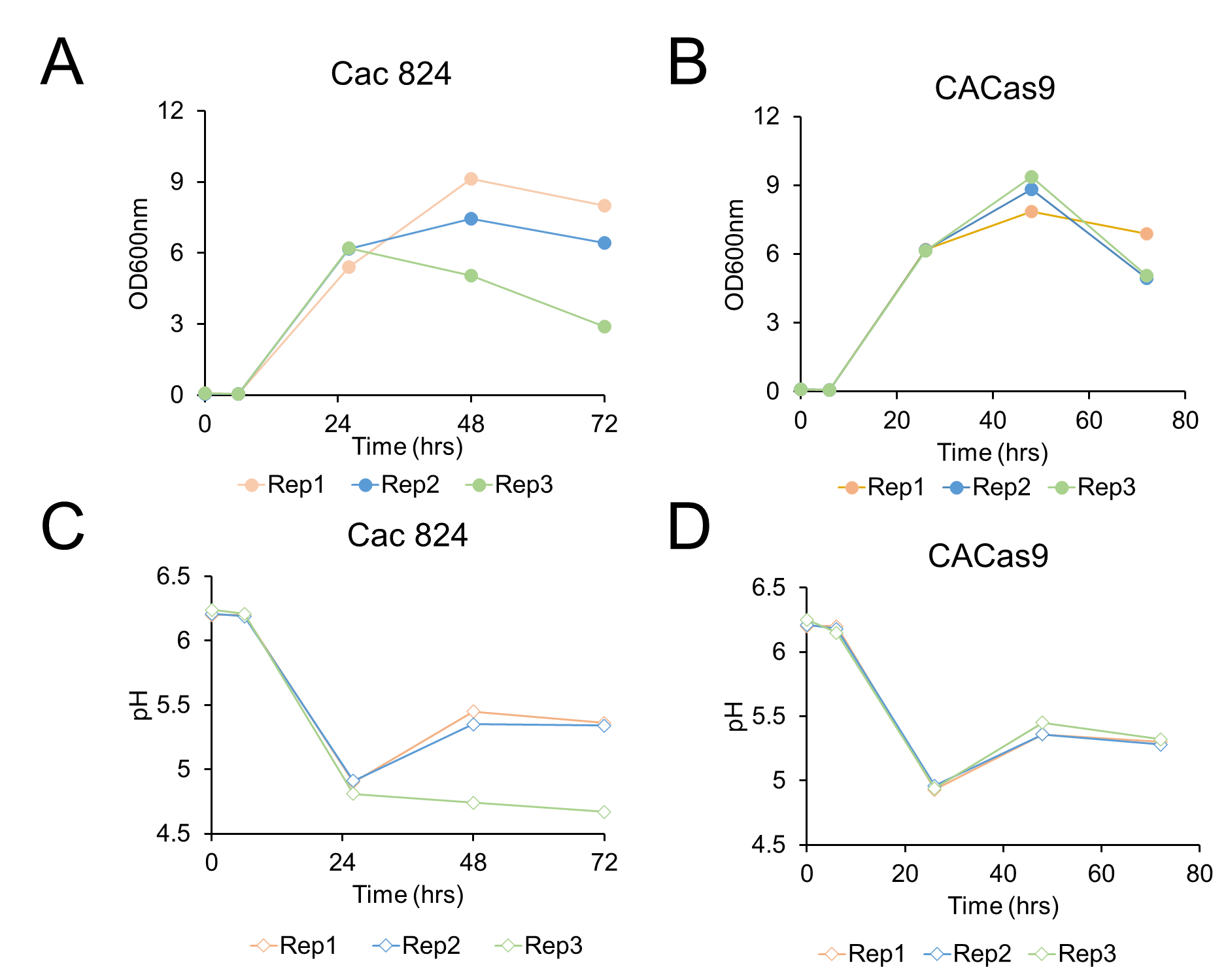
**Supplementary Figure 2.** Effects of butyrate on acetate assimilation, glucose consumption, and growth of CACas9 ∆*hbd.* (A) Profiles of acetate at different initial butyrate concentrations. (B) Profiles of glucose consumption at different initial butyrate concentrations. (C) Growth kinetics and specific growth rates in CGM and CGMB media. The data represent mean ± standard deviation from three biological replicates.

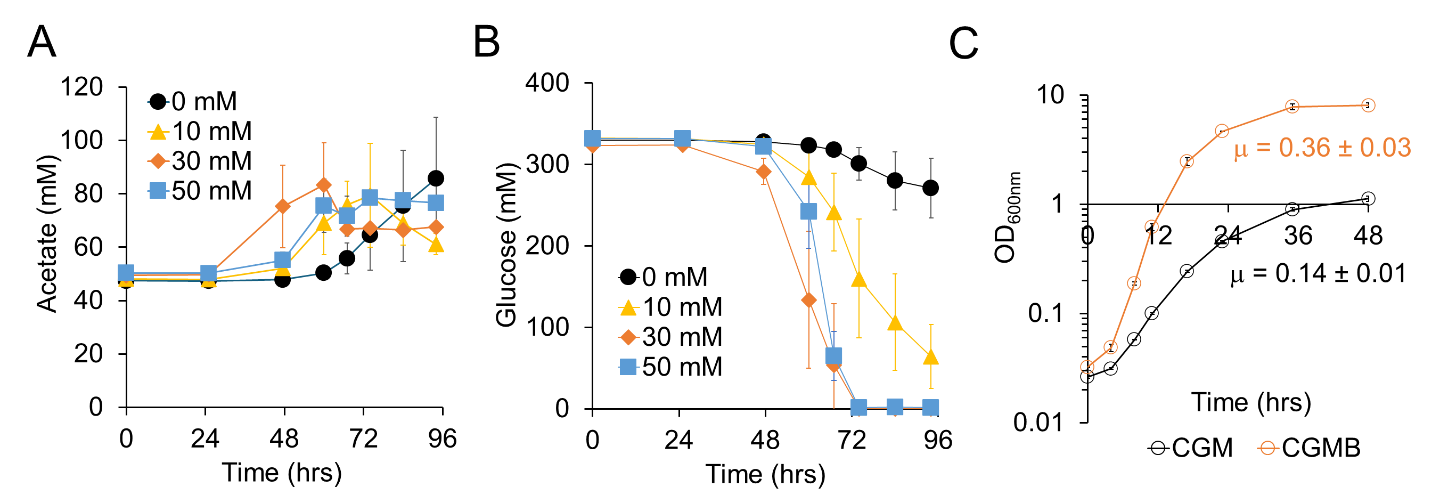

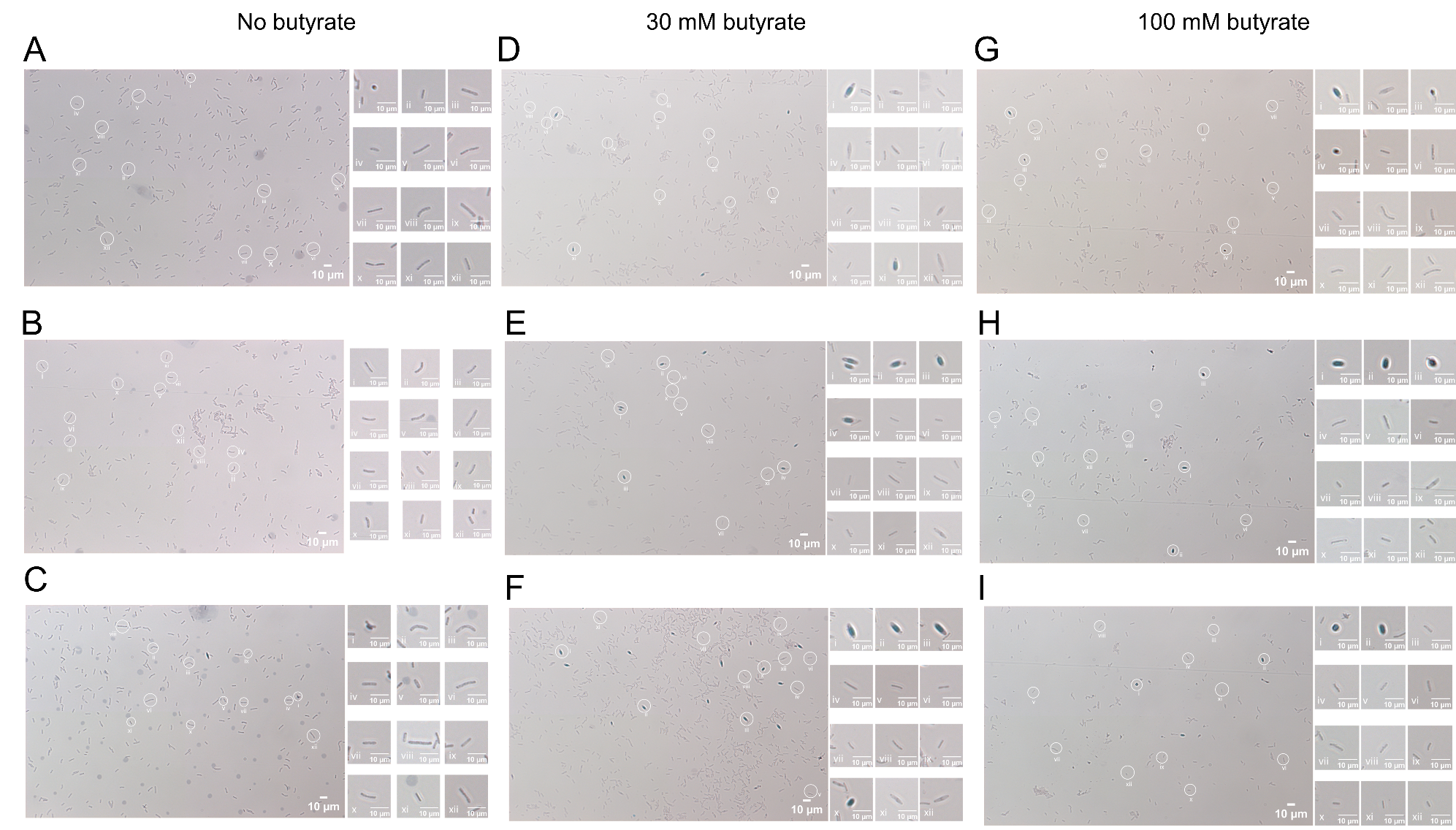
 **Supplementary Figure 3.** Effects of butyrate on sporulation of CACas9 ∆*hbd.* (A-C). Microscopic images of cells from cultures without butyrate addition. (D-F) Microscopic images of cells from cultures with 30 mM butyrate addition. (G-I) Microscopic images of cells from cultures with 100 mM butyrate addition.

**Supplementary Figure 4.** Glucose consumption and acetone production by CACas9 ∆*hbd* ∆*spo0A* in CGM and CGMB media. (A) Profiles of glucose consumption. (B) Profiles of acetone production. All the data represent mean ± standard deviation from three biological replicates.

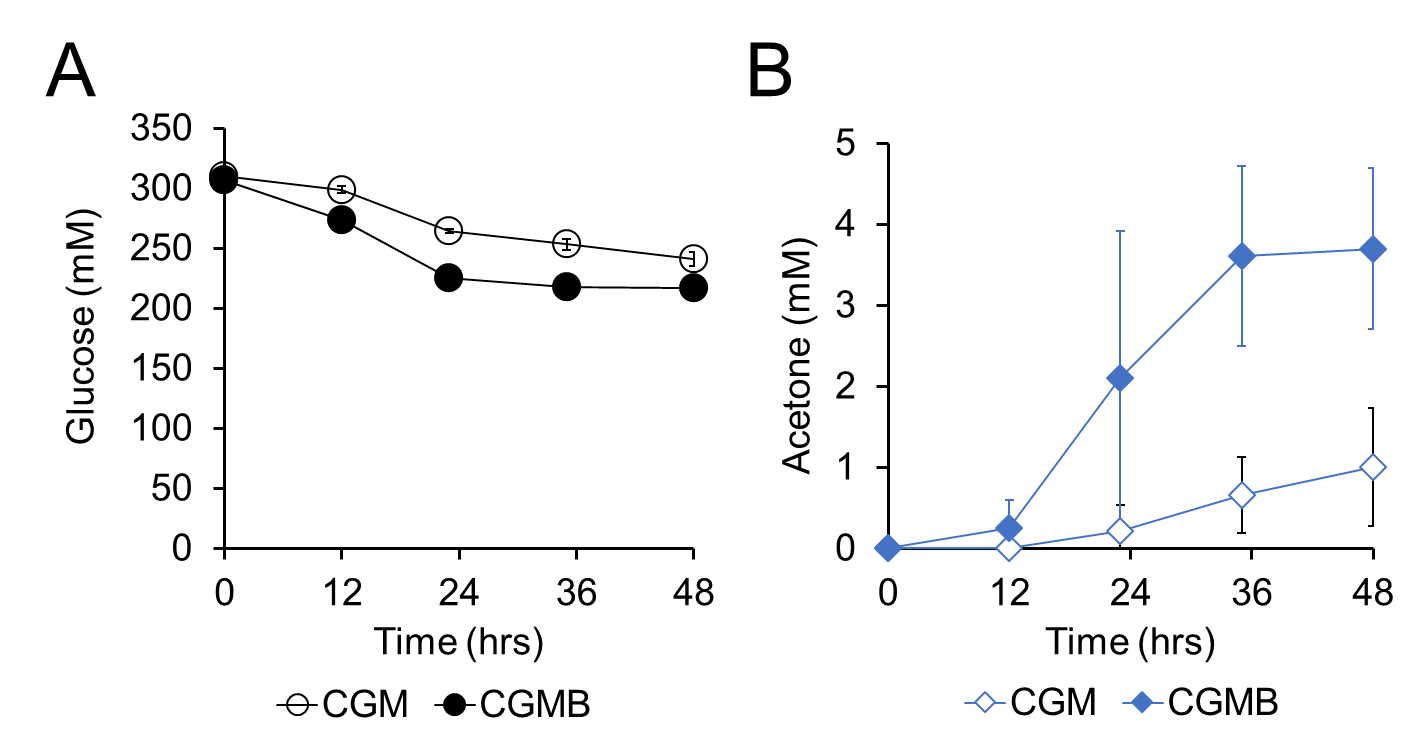

**Supplementary Figure 5.** Integration of spCas9 in the chromosome of *C. acetobutylicum* M5. (A) Structure of chromosomal integration location. Amplification of the location from a parental M5 strain results in 1.9kb of PCR product, while 7.9kb of PCR product is expected from a Cas9 integrated M5 strain. (B) An agarose gel picture of colony PCR products from seven transformant colonies and a parental M5 strain. The numbers above each well indicate individual transformant colonies. M, DNA ladder; C, parental M5. The red arrow indicates the expected size (7.9kb) of the integrated Cas9 cassette.

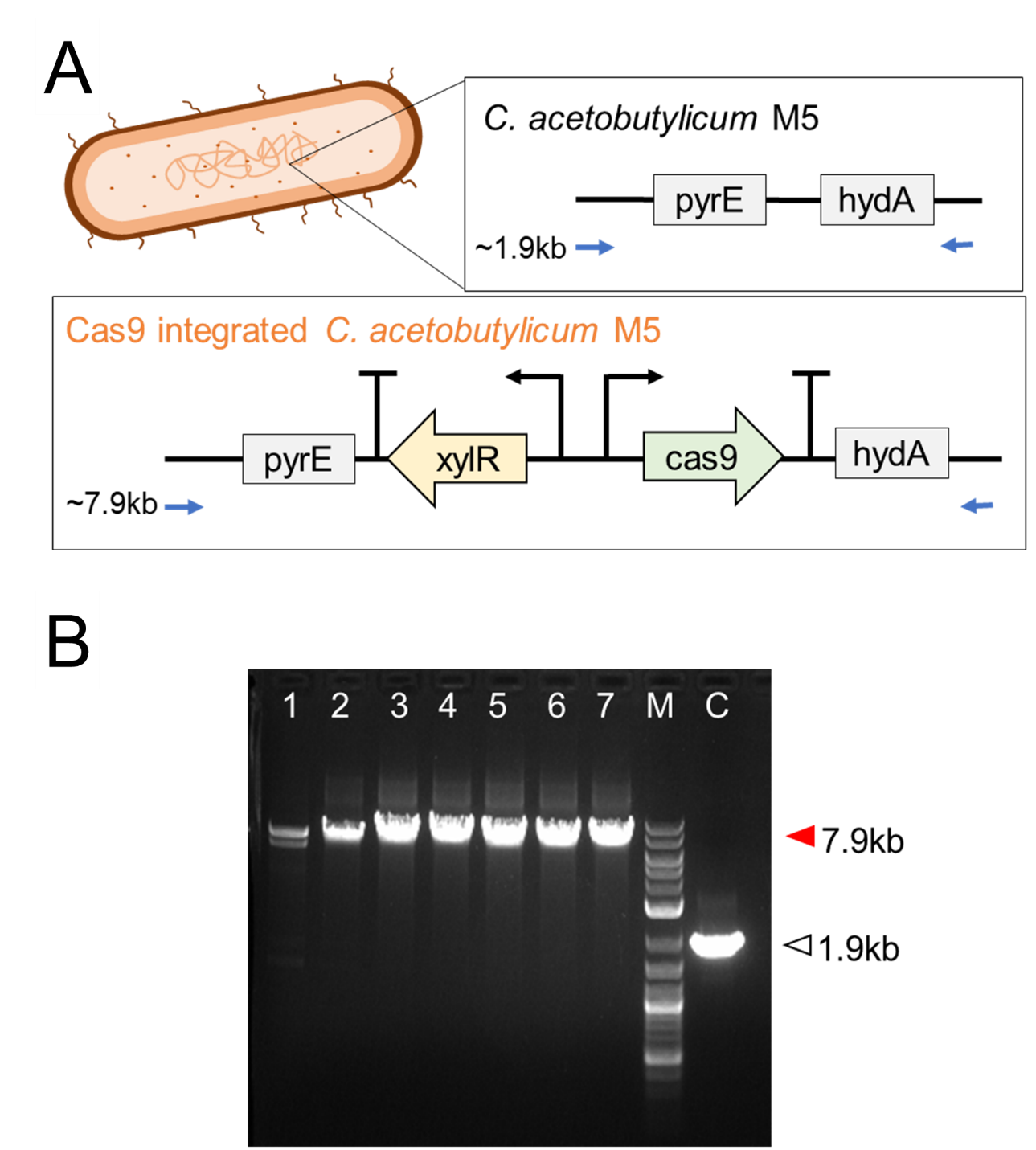

**Supplementary Figure 6.** Adaptive laboratory evolution of M5Cas9 ∆*hbd*::*frt-cat-frt* in the absence of butyrate. The data represent mean ± standard deviation from two biological replicates.

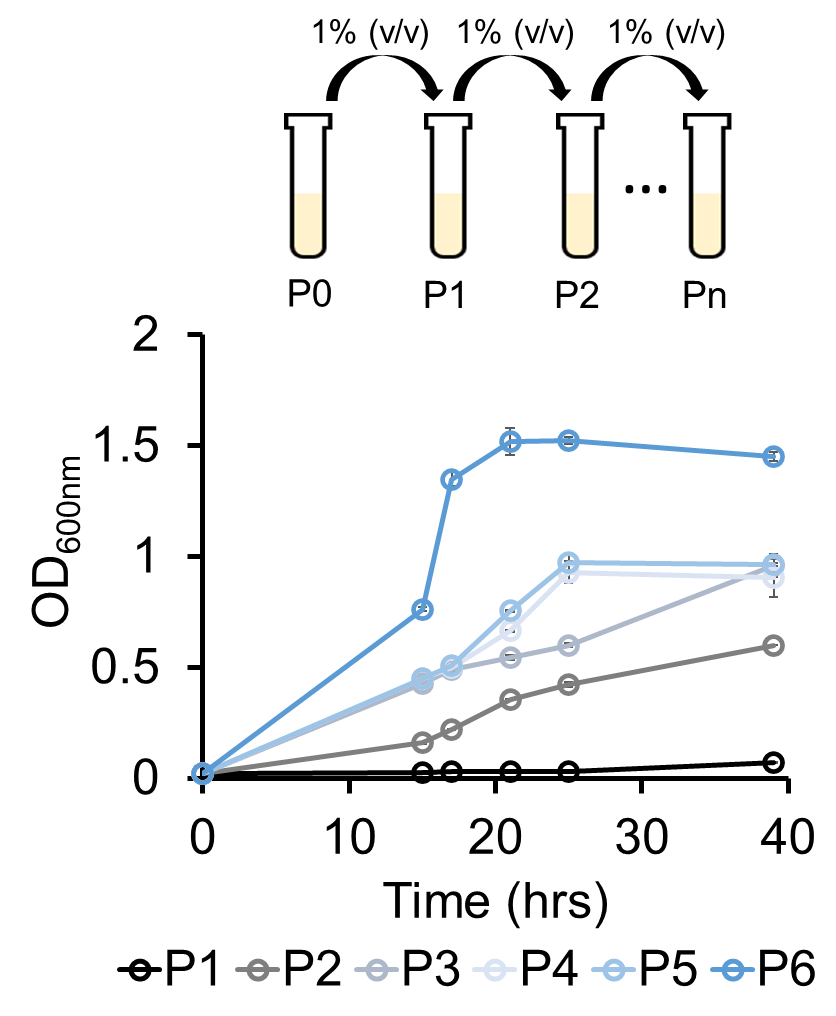

**Supplementary Figure 7.** Growth of M5Cas9 ∆*hbd* with and without crotonate addition. The data represent mean ± standard deviation from two biological replicates.

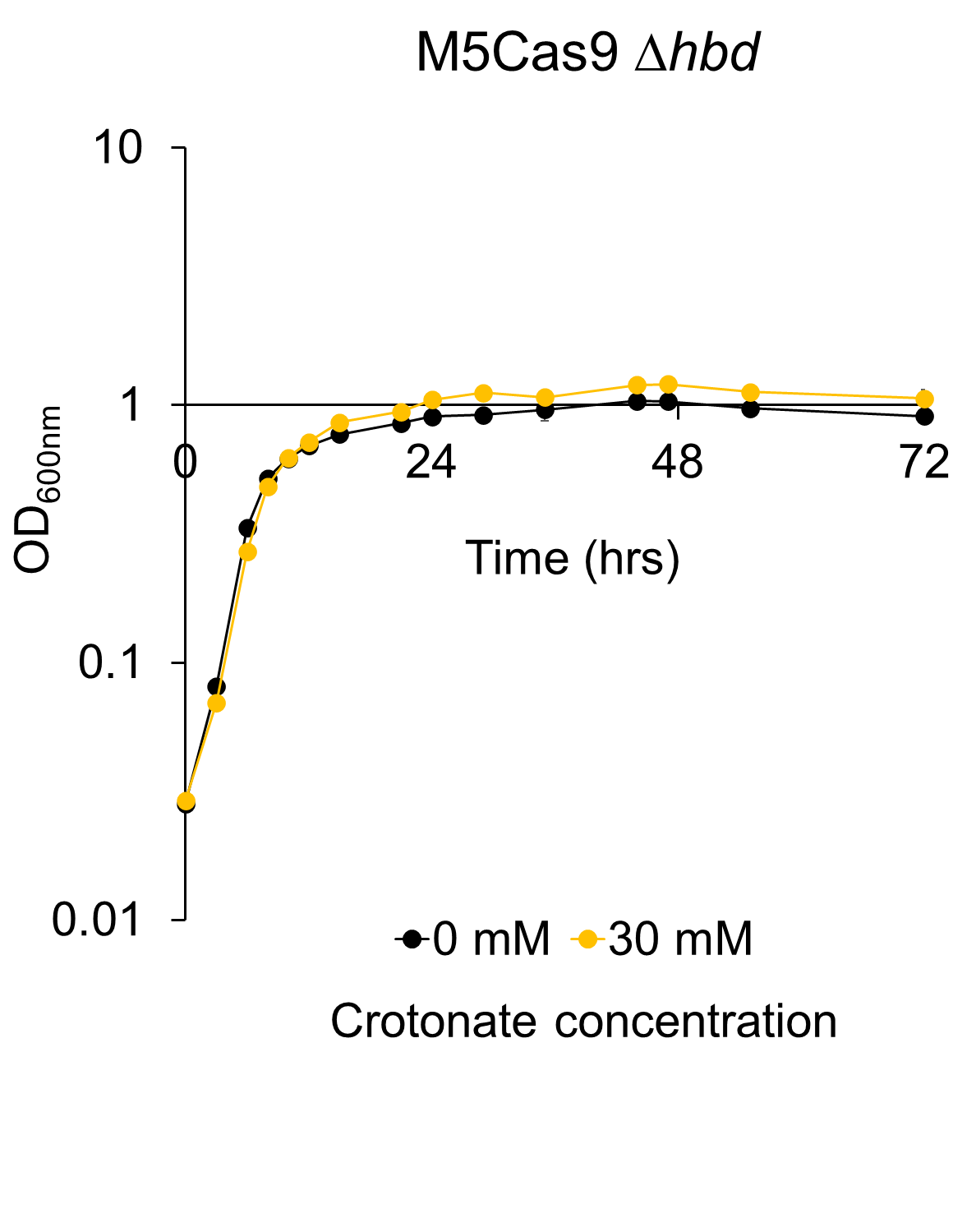

**Supplementary Figure 8.** Effects of butyrate on the cell growth of non-butyrate forming bacteria. (A) Growth of *C. ljungdahlii* without or with 30 mM butyrate. (B) Specific growth rates of *C. ljungdahlii* without or with 30 mM butyrate. (C) Growth of *E. coli* without or with 30 mM butyrate. (D) Specific growth rates of *E. coli* without or with 30 mM butyrate. All the data represent means ± 1 standard deviation from three biological replicates. N.S., statistically not significant (p-value > 0.05); *, p-value < 0.01 from a two tailed Student’s t-test.

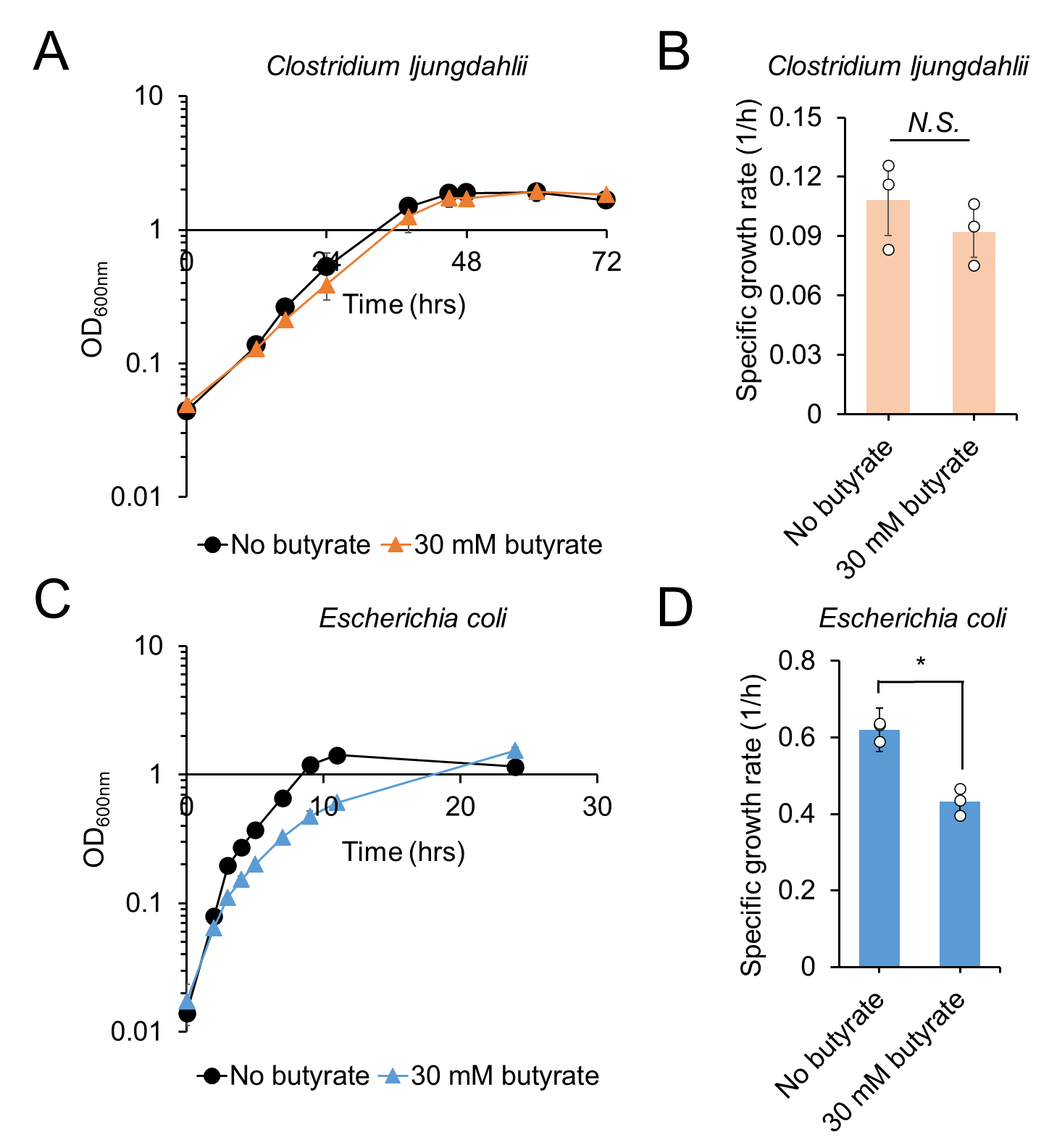
